# Robust Estimation of the Phylogenetic Origin of Plastids Using a tRNA-Based Phyloclassifier

**DOI:** 10.1101/442608

**Authors:** Travis J. Lawrence, Katherine C. H. Amrine, Wesley D. Swingley, David H. Ardell

## Abstract

The trait of oxygenic photosynthesis was acquired by the last common ancestor of Archaeplastida through endosymbiosis of the cyanobacterial progenitor of modern-day plastids. Although a single origin of plastids by endosymbiosis is broadly supported, recent phylogenomic studies report contradictory evidence that plastids branch either early or late within the cyanobacterial Tree of Life. Here we describe CYANO-MLP, a general-purpose phyloclassifier of cyanobacterial genomes implemented using a Multi-Layer Perceptron. CYANO-MLP exploits consistent phylogenetic signals in bioinformatically estimated structure-function maps of tRNAs. CYANO-MLP accurately classifies cyanobacterial genomes into one of eight well-supported cyanobacterial clades in a manner that is robust to missing data, unbalanced data and variation in model specification. CYANO-MLP supports a late-branching origin of plastids: we classify 99.32% of 440 plastid genomes into one of two late-branching cyanobacterial clades with strong statistical support, and confidently assign 98.41% of plastid genomes to one late-branching clade containing unicellular starch-producing marine/freshwater diazotrophic Cyanobacteria. CYANO-MLP correctly classifies the chromatophore of *Paulinella chromatophora* and rejects a sister relationship between plastids and the early-branching cyanobacterium *Gloeomargarita lithophora*. We show that recently applied phylogenetic models and character recoding strategies fit cyanobacterial/plastid phylogenomic datasets poorly, because of heterogeneity both in substitution processes over sites and compositions over lineages.

## Introduction

The acquisition of a cyanobacterial endosymbiont by the last common ancestor of Archaeplastida [1, 36, 38] transferred the trait of oxygenic photosynthesis to eukaryotes over one billion years ago [20]. The diversity of eukaryotic photoautotrophs radiating from this event profoundly transformed the terrestrial biosphere through changes to primary biomass production, atmospheric oxygenation, and the colonization of new ecosystems [26].

It is widely accepted both that the plastids originated in a single primary endosymbiotic event [37], and that the photosynthetic chromatophore of the freshwater amoeba *Paulinella chromatophora* evolved later in a second primary endosymbiotic event [25, 40]. However, despite substantial progress on a robust cyanobacterial Tree of Life (CyanoToL) [9, 39, 45, 47], the root of plastids within the CyanoToL remains controversial. Recent phylogenomic studies strongly support contradictory conclusions, with plastids branching either early [12, 43, 47, 52] or late [8, 13, 20, 39] within the CyanoToL. In contrast, orthogonal evidence from endosymbiotic gene transfers [16] and eukaryotic evolution of glycogen and starch metabolic pathways [6, 15] consistently support a late-branching origin of plastids within the CyanoToL.

Phylogenetic inferences concerning plastid origins are complicated by large evolutionary distances accumulated over at least one billion years of vertical descent, by extreme reduction of genomes in plastids [53] and Cyanobacteria [18, 44], and by secondary and tertiary endosymbiotic acquisitions of plastids. Furthermore, reductive genome evolution alters the stationary nucleotide composition of genomes and gene products [7], violating the assumptions and applicability of many phylogenetic models [10, 17, 21, 28, 42].

Recently, we introduced a machine learning approach to the phyloclassification of genomes based on scoring tRNA gene complements against bioinformatically estimated taxon-specific tRNA functional signatures called tRNA Class-Informative Features (CIFs) [3]. tRNA CIFs, as visualized in function logos [22], contain information [46] about the functional identity of tRNAs for tRNA-interacting proteins. We demonstrated the strong recall and accuracy of a tRNA-CIF-based alpha-proteobacterial phyloclassifier despite convergent non-stationary compositions of alpha-proteobacterial tRNA genes, and likely horizontal transfers of genes for tRNAs and tRNA-interacting proteins [3].

In the present work, we improved our tRNA-based phyloclassifier approach and applied it to investigate the origin of plastids within the CyanoToL. Based on 5,270 tRNA gene sequences from 113 cyanobacterial genomes, our CYANO-Multi-Layer Perceptron (CYANO-MLP) phyloclassifier consistently classifies 433 plastid genomes within the B2 and B3 sister clades of Cyanobacteria [47]. These clades include marine/freshwater unicellular diazotrophic species previously noted to share synapomorphic starch metabolic pathway traits with plastids [15, 20]. We reconciled our results with prior work by demonstrating that recently applied phylogenetic models and character recoding strategies fit cyanobacterial/plastid phylogenomic datasets poorly because of heterogeneity of substitution processes over sites and lineage-specific compositional biases.

## Materials and Methods

### tRNA Gene Data and Genome Sets

From NCBI, we downloaded the set *S* of 117 cyanobacterial genomes analyzed in [47], the set *Gl* of one genome of the cyanobacterium *Gloeomargarita lithophora*, the set *Pc* of one genome of the chromatophore of the fresh-water amoeba *Paulinella chromatophora*, and the set *P* of 440 complete plastid genomes containing representatives from all three lineages of Archaeplastida (Glaucocystophyta, Rhodophyta, and Viridiplantae). Let *C ≡ S* ∪ *Gl* ∪ *Pc*. For every genome *g* ∈ *C*, we annotated a set *T_g_* of tRNA genes as the union of predictions from tRNAscan-SE v1.31 [34] in bacterial mode and ARAGORN v1.2.36 [31] with default settings. We annotated tRNA genes in the set *P* of plastid genomes similarly, except we discarded as false positives gene predictions from ARAGORN that contained introns in tRNA isotypes that have not been previously described to contain introns [35, 48, 54]. We additionally filtered away tRNA gene predictions for land plant plastid genomes that contained anticodons not previously observed in land plant plastid tRNA genes [2, 50].

**Figure 1.**
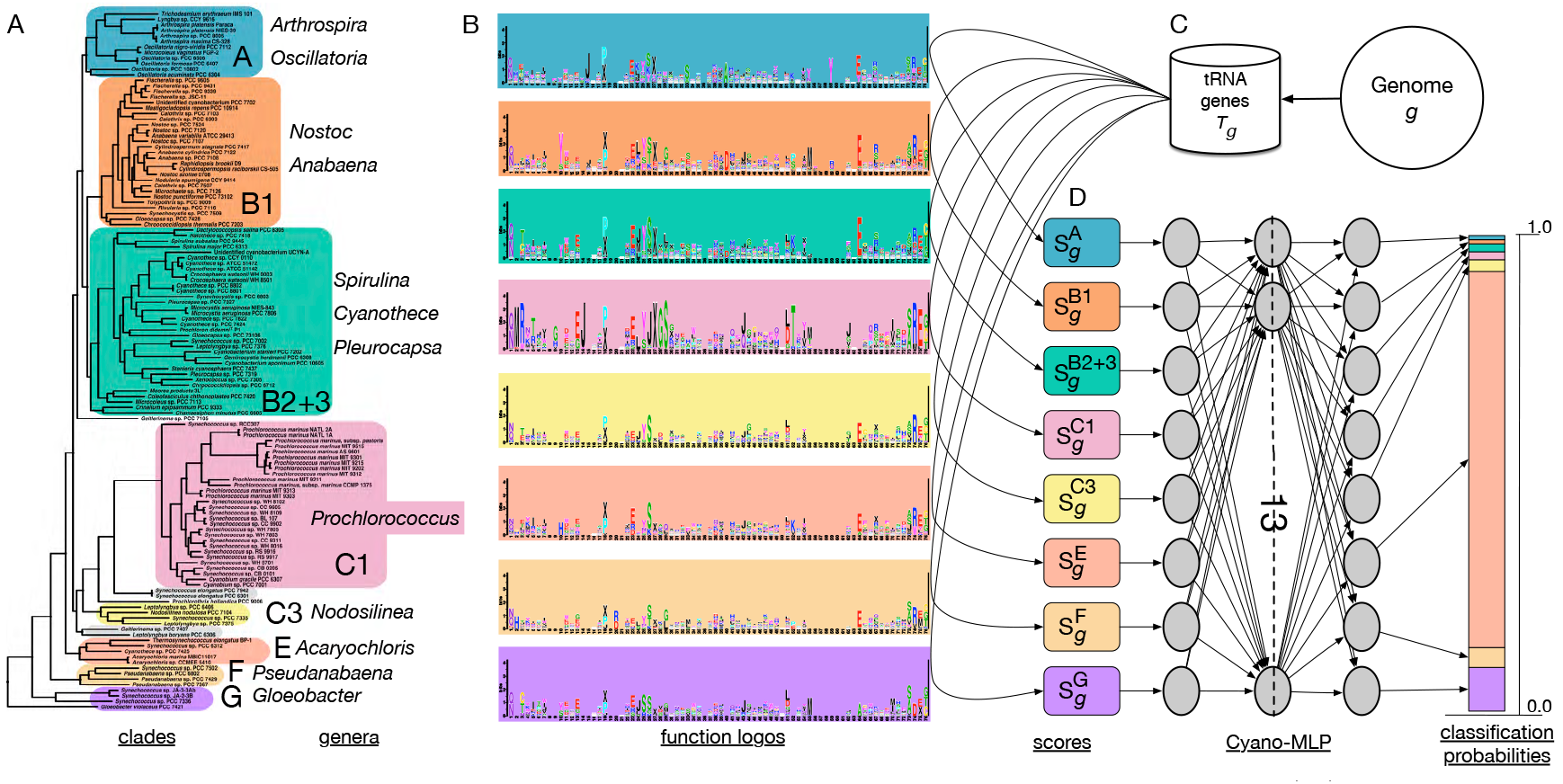
Schematic overview of CYANO-MLP phyloclassification workflow. (A) The cyanobacterial phylogeny from [47] was used to define cyanobacterial clades for which we estimated tRNA CIFs. Cyanobacterial clades, indicated by background colors, were named according to [47], except for clade B2+3, which combines clades B2 and B3. Clades with grey backgrounds were excluded from analysis because of limited genome sample sizes. Next, tRNA genes are predicted from cyanobacterial genomes and combined by clade for function logo estimation [22]. (B) Example function logos, for Uracil, with background colors corresponding to each cyanobacterial clade. (C) For a genome *g* to be classified, tRNA gene complements *T_g_* are predicted and scored against the logos for each clade, to produce input score vectors for CYANO-MLP training and classifications. (D) The architecture of the artificial neural network CYANO-MLP, and a classification probability vector output from CYANO-MLP represented as a stacked bar chart.

We annotated the functional types of tRNA genes either as elongator isotypes by anticodon alone or, for those containing the CAU anticodon, into initiator tRNA ^Met^ (”X”), elongator tRNA ^Met^, or tRNA ^Ile^ _CA U_ (”J”) using TFAM v1.4 [4] with the TFAM model used in [3, 5]. We aligned tRNA sequences using COVEA v2.4.4 [19] and the prokaryotic tRNA covariance model from tRNAscan-SE [34]. We edited the alignment by first removing sites containing 99% or more gaps using FAST v1.6 [32], and then removing sequences with unusual secondary structure. Lastly, we mapped sites to Sprinzl coordinates [49] and manually removed the variable arm, CCA tail, and sites not mapping to a Sprinzl coordinate using Seaview v4.6.1 [24]. The alignment is available as supplementary data.

We partitioned cyanobacterial tRNA genes into sets *T_g_* for each genome *g* of origin, and separately into sets *T_X_* for each cyanobacterial clade *X*, with *X* ∈ *CC* ≡ {*A, B*1, *B*2 + 3, *C*1, *C*3, *E, F, G*} corresponding to clades identified in [47], except for fusion of clades B2 and B3 into their union B2+3 and exclusion of four genomes in two clades, C2 and D, for insufficient data as defined by yielding fewer than 120 tRNA genes (Fig. 1). Let *R* ⊂ *S* be the set of all 113 cyanobacterial genomes not excluded. For every genome *g* ∈ *R* and each clade *X* ∈ *CC*, we also created leave-one-out cross-validation training sets 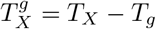.

### Genome Scoring

Following Amrine et al. [3], we produced training input vectors by first calculating clade-dependent Gorodkin heights [3, 23] 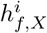, in function logos [22] for all clade-specific tRNA gene sets *T_X_* or 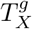 with *X* ∈ *CC*, for all features *f* ∈ *F* ≡ {*A, C, G, U* } × *SC*, where *SC* is the set of Sprinzl Coordinates [49], and for all functional types *i* ∈ *I* ≡ *A* ⋃ {*J, X*}, where *A* is the set of short IUPAC amino acid symbols standing for aminoacylation identities of elongators. We computed function logos using custom software TSFM available at https://github.com/tlawrence3/tsfm/tree/v0.9.6.

To score the tRNA gene complement *T_g_* of genome *g*, we calculated a vector of tRNA CIF-based scores 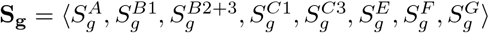, in which element 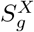 is the average, over all genes *t* ∈ *T_g_* of any type *i_t_* ∈ *I*, where *i_t_* is the type of gene *t*, of the sum over all features *f* ∈ *t* ⊂ *F* contained in that gene, of the Gorodkin heights [23] of those features for genes of that type in clade *X* ∈ *CC*:

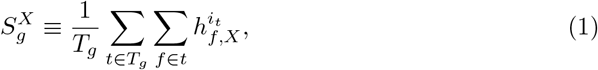

Following recommended practice [11], we standardized score vectors of both training and query data by subtracting the mean score vector of training data and dividing element-wise by the standard deviations of scores by clade. Let 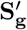 be the standardized score vector of **S_g_**.

### Phyloclassifier Model Training and Optimization

We implemented our multilayer neural network phyloclassifier using the MLPClassifier API of scikit-learn v0.18.1 [41] in Python v3.5.2. We trained models for up to 2000 training epochs, stopping early if for two consecutive iterations the Cross-Entropy loss function value did not decrease by a minimum of 1 × 10^*−*4^, and with random shuffling of data between epochs. We used the rectifier activation function for hidden layer neurons, the L-BFGS algorithm for weight optimization, and an alpha value of 0.01 for the L2 regularization penalty parameter. Lastly, we used the soft-max function to calculate classification probability vectors. Using leave-one-out cross-validation (LOOCV), we optimized neural network architecture for accuracy averaged over genomes *g∈ R* considering all architectures with from one to four hidden layers and each layer individually containing from eight to sixteen nodes. To test the statistical significance of the average accuracy from LOOCV of the architecture-optimized CYANO-MLP, we permuted clade labels over training data in 100,000 replicates, followed by LOOCV and model retraining for each replicate.

### Phyloclassification and Bootstrapping

For each genomic tRNA gene set *T_g_*, with *g P*⋃ *Pc* ⋃ *Gl*, we computed a standardized score vector 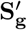, input this to CYANO-MLP, and classified to the clade with largest classification probability. To examine the consistency of phylogenetic signals in our data, we computed 100 bootstrap replicates of sites in our alignment of training and test tRNA gene data, followed by CIF-estimation, model retraining, and genome scoring and classification with each bootstrap replicate of CYANO-MLP. We summarized bootstrap results for cyanobacterial genomes by the number of replicates in which the most probable classification for a genome was its true clade of origin.

### Leave-Clade-Out and Balanced Model Variants

To examine the sensitivity of CYANO-MLP to missing data and model mis-specification, we re-optimized and re-trained models after leaving out one cyanobacterial clade or using only cyanobacterial clades A, B1, and B2+3. To produce clade-balanced training datasets, we randomly resampled training score vectors so that each cyanobacterial clade had sample sizes equal to the best-sampled clade, and then re-optimized and re-trained models.

### Evaluation of Phylogenetic Model Adequacy

We examined goodness of fits of the phylogenomic datasets of Shih *et al.* [47], Ponce-Toledo *et al.* (chloroplast-marker dataset) [43] and Ochoa de Alda *et al.* (dataset 11) [39] with the substitution models originally used in those studies, namely LG+4Γ [33] and CAT-GTR+4Γ [28, 29]. Posterior Predictive Analyses (PPA) were performed to test fits for site-specific constraint biases using PPA-DIV [28] and across-lineage compositional biases using PPA-MAX and PPA-MEAN [10]. Additionally, we assessed model adequacy under three amino acid recoding strategies, Dayhoff-6 (Day6) [14], the six-state recoding strategy of Susko and Roger (SR6) [51], and the six-state recoding strategy of Kosiol et al. (KGB6) [27]. PPA results were interpreted using Z-scores under the assumption that the test statistics follow a normal distribution. We used a Z-score threshold of *Z* ≥ 5 as strong evidence for rejecting the model. We performed phylogenetic analyses using Phylobayes MPI v1.8 [30] with at least 1000 replicates and running two MCMC chains in parallel for each analysis. Convergence of chain trajectories was assessed using TRACECOMP and BBCOMP utilities provided with Phylobayes MPI. Convergence was assumed when the discrepancies of model parameters and bipartition frequencies between independent chains was less than 0.18. The number of cycles to discard as burn-in was determined by visually examining the traces of the log-likelihood and other model parameters for stationarity using Tracer v1.6.0.

## Results

### tRNA Data and CIF estimation

We annotated and extracted 5,476 tRNA genes from the 117 cyanobacterial genomes analyzed in [47], averaging 46.80 tRNA genes per cyanobacterial genome, 14,841 tRNA genes in 440 Archaeplastida plastid genomes averaging 33.73 tRNA genes per plastid genome, 44 tRNA genes from the Cyanobacterium *Gloeomargarita lithophora*, and 42 tRNA genes from the chromatophore genome of the fresh-water amoeba *P. chromatophora* (Table 1; Supplemental File 1). We excluded four genomes from further analysis (Fig. 1) and estimated function logos for cyanobacterial clades A, B1, B2+3, C1, C3, E, F, and G (Fig. 1, S1-S8; Table S1,S2) using the clade nomenclature of [47]. We fused clades B2 and B3 because they are sister clades and B3 contained only one genome. The C1 clade had the biggest sample with a divergent nucleotide composition, elevated in contents of G and C (Table 1). Interestingly, clade C1 exhibited many gains of Uracil CIFs (Fig. 1B) and also Adenine CIFs (Fig. S4).

**Table 1.** Summary statistics on genomes and tRNA genes of cyanobacterial and plastid clades and grades.

| Clade/Grade | Genomes (G) | Genes (T) | T/G | Bases (N) | N/T | %A | %T | %G | %C |
| --- | --- | --- | --- | --- | --- | --- | --- | --- | --- |
| **Cyanobacteria** |  |  |  |  |  |  |  |  |  |
| A | 11 | 555 | 50.45 | 40,362 | 72.72 | 20.1 | 23.3 | 31.2 | 25.4 |
| B1 | 27 | 1,395 | 51.67 | 101,640 | 72.86 | 19.7 | 23.3 | 31.6 | 25.4 |
| B2+3 | 30 | 1,314 | 43.80 | 95,685 | 72.82 | 19.5 | 23.0 | 32.0 | 25.5 |
| C1 | 29 | 1,205 | 41.55 | 87,856 | 72.91 | 18.8 | 21.8 | 32.7 | 26.7 |
| C2 | 2 | 90 | 45.00 | 6,550 | 72.78 | 19.5 | 21.9 | 32.2 | 26.4 |
| C3 | 3 | 142 | 47.33 | 10,346 | 72.86 | 19.1 | 22.8 | 32.3 | 25.7 |
| D | 2 | 116 | 58.00 | 8,430 | 72.67 | 19.7 | 23.3 | 31.7 | 25.3 |
| E | 5 | 266 | 53.20 | 19,378 | 72.85 | 19.3 | 22.8 | 32.1 | 25.8 |
| F | 4 | 211 | 52.75 | 15,339 | 72.70 | 20.1 | 23.4 | 31.3 | 25.2 |
| G | 4 | 182 | 45.50 | 13,218 | 72.63 | 18.7 | 21.7 | 32.8 | 26.8 |
| *G. lithophora* | 1 | 44 | 44 | 3,194 | 72.59 | 18.8 | 22.7 | 32.3 | 26.2 |
| **Plastids/Chromatophores** |  |  |  |  |  |  |  |  |  |
| Charophyta | 10 | 352 | 35.20 | 25,513 | 72.48 | 21.1 | 24.2 | 30.1 | 24.6 |
| Chlorophyta | 7 | 226 | 32.29 | 16,436 | 72.73 | 21.9 | 25.4 | 29.1 | 23.6 |
| Cryptophyta | 4 | 117 | 29.25 | 8,512 | 72.75 | 20.9 | 24.2 | 29.9 | 25.0 |
| Heterokonta | 33 | 992 | 30.06 | 72,229 | 72.81 | 21.2 | 25.1 | 29.6 | 24.1 |
| Eudicots | 191 | 6,593 | 34.52 | 478,337 | 72.55 | 21.4 | 25.2 | 29.7 | 23.7 |
| Euglenaceae | 10 | 276 | 27.60 | 20,070 | 72.72 | 22.3 | 27.1 | 28.4 | 22.3 |
| Monilophytes | 8 | 270 | 33.75 | 19,641 | 72.74 | 21.2 | 24.5 | 30.0 | 24.3 |
| Gymnospermae | 26 | 824 | 31.69 | 59,867 | 72.65 | 21.9 | 24.6 | 29.6 | 23.9 |
| Haptophyta | 4 | 111 | 27.75 | 8,079 | 72.78 | 21.0 | 25.1 | 29.7 | 24.2 |
| Monocots | 112 | 3930 | 35.10 | 285,302 | 72.60 | 21.7 | 25.2 | 29.5 | 23.7 |
| Magnoliids | 9 | 315 | 35.00 | 22,856 | 72.56 | 21.3 | 24.9 | 29.7 | 24.0 |
| Nymphaeales | 2 | 68 | 34.00 | 4,934 | 72.56 | 21.4 | 25.1 | 29.8 | 23.7 |
| Rhodophyta | 20 | 624 | 31.20 | 45,438 | 72.82 | 21.9 | 25.2 | 29.1 | 23.8 |
| Bryophyta | 3 | 108 | 36.00 | 7,845 | 72.64 | 22.2 | 25.4 | 29.0 | 23.4 |
| Glaucocystophyta | 1 | 35 | 35 | 2,543 | 72.66 | 20.4 | 23.7 | 30.9 | 25.0 |
| *P. chromatophora* | 1 | 42 | 42 | 3,060 | 72.86 | 19.1 | 21.7 | 32.7 | 26.5 |

### Training and Validation of a tRNA-Based Cyanobacterial Phyloclassifier

We trained CYANO-MLP on input vectors generated by scoring cyanobacterial genomic tRNA gene complements against clade-specific cyanobacterial CIFs as described in Methods. We systematically optimized the parameters and architecture of CYANO-MLP on the training data, settling on a single hidden layer of 13 nodes (Fig. 1), which achieved an average accuracy of 0.8673 (permutation test; *p* = 0.0001), calculated using Leave-One-Out Cross-Validation (LOOCV; Fig. S9, S10; Table S3). To examine the effects of unbalanced training data on the performance of CYANO-MLP, we created a separate clade-balanced version of the model (CYANO-MLP-BAL) by resampling data from under-represented clades. CYANO-MLP-BAL achieved a LOOCV average accuracy of 0.9875 with improvements in precision and recall for all clades (fig. S19; Table S3), suggesting that biased sampling is an important and addressable limitation to the accuracy of CYANO-MLP. In addition, cyanobacterial reclassifications were correct in at least 97 of 100 bootstrap replicates of CYANO-MLP, showing that phylogenetic signals are consistent across tRNA CIFs (Supplemental File 1).

A desirable attribute of a phyloclassifier is an ability to signal “none-of-the-above” when the true clade of a query is unrepresented in the model. To address robustness to model specification and investigate the ability of CYANO-MLP to signal “none-of-the-above”, we trained additional versions of CYANO-MLP that leave out the largest-sampled clades, namely A (CYANO-MLP[!A]), B1 (CYANO-MLP[!B1]), B2+3 (CYANO-MLP[!B2+3]), or C1 (CYANO-MLP[!C1]). For each model variant, we then reclassified all genomes, including from the clade that had been left out. Overall, average LOOCV accuracies were similar to CYANO-MLP for each classifier variant (Fig. S14-S18; Table S3) with CYANO-MLP[!A] having the largest gain in accuracy (LOOCV: 0.9216; Fig. S16; Table S3) over CYANO-MLP. This was not unexpected, given that clade A had the lowest precision (Fig. S15-S18) and the smallest sample size among left-out clades (Table 1). Furthermore, the recall of clade A was most improved in CYANO-MLP-BAL. Generally with CYANO-MLP, classifications of cyanobacterial genomes from excluded clades were more equivocal than those of genomes from represented clades (Table S9-S12; Supplemental File 1). We claim that equivocal classifications with CYANO-MLP signal “none-of-the-above.”

### The *Paulinella chromatophora* Chromatophore Phyloclassifies to the Marine C1 *Prochlorococcus*/*Synechococcus* Clade

The phylogenetic origin of the *P. chromatophora* chromatophore from marine *Prochlorococcus*/*Synechococcus* clade (clade C1; Fig. 1) is well-supported by several phylogenomic analyses [39, 47, 52]. CYANO-MLP classified the *P. chromatophora* chromatophore to clade C1 concordantly with a 99.98% probability and 100% bootstrap support (Fig. 2; Table S5). Additionally, this phyloclassification was robust to model specification and obtained also with CYANO-MLP-BAL, CYANO-MLP[!A]), CYANO-MLP[!B1], and CYANO-MLP[!B2+3]. Finally, the *P. chromatophora* chromatophore classified similarly to other C1 genomes when using CYANO-MLP[!C1] (Table S11,S13).

**Figure 2.**
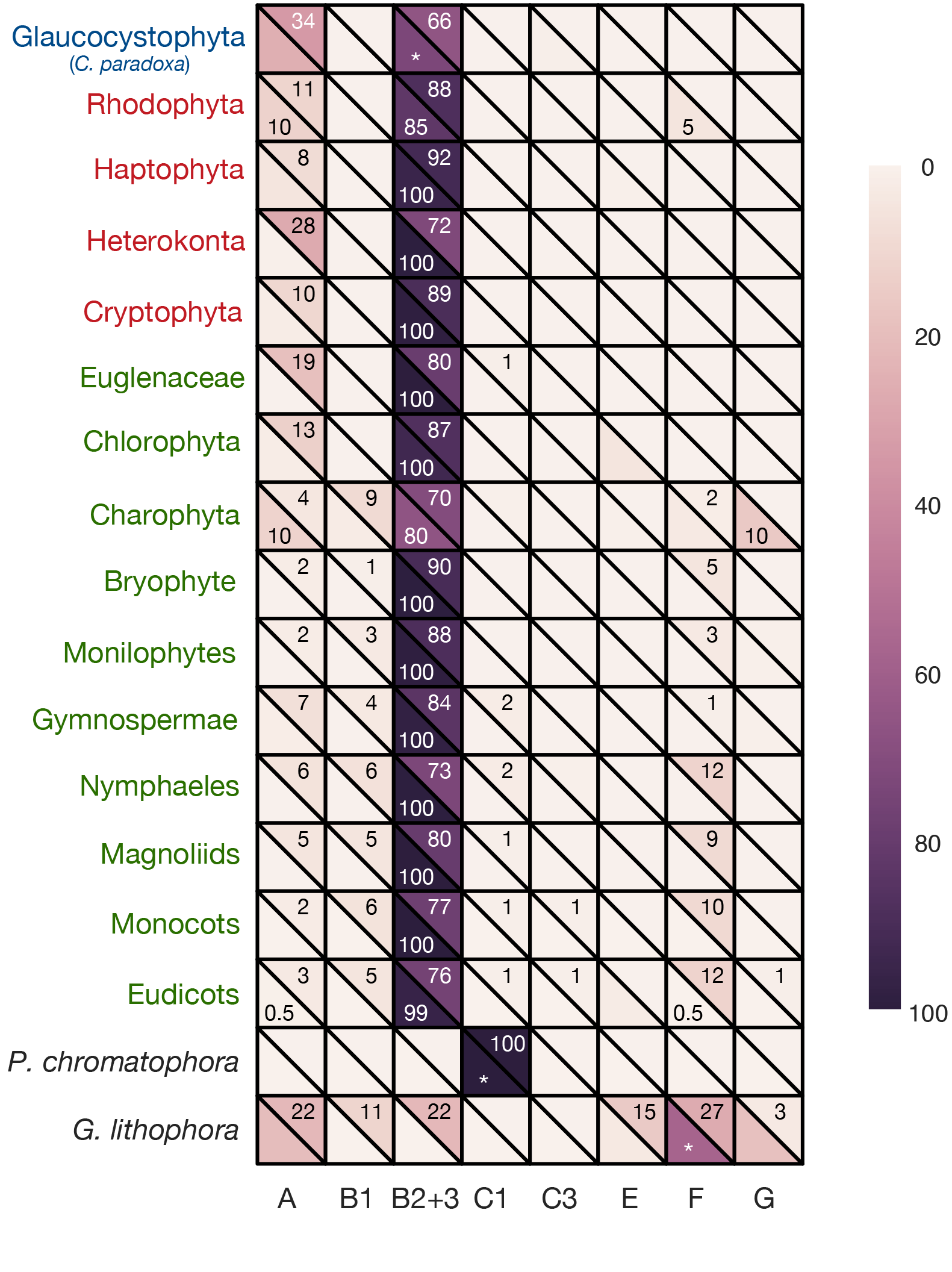
CYANO-MLP classification results for genomes of plastids, the chromatophore of *P. chromatophora*, and the cyanobacterium *G. lithophora*. Row label colors denote Archeaplastid clades with Rhodophyta in red, Chloroplastida in green, and Glaucocystophyta in blue, or non-Archaeplastida in black. Heatmap lower half-cells show average probabilities of classifications of genomes to clades with text labels denoting percentages of genomes classifying to cyanobacterial clades. Asterisks (*) denote single genomes. Heatmap upper half-cell colors (and text labels) show median bootstrap classification frequencies (as percentages). Absence of labels denotes zero frequencies.

### CYANO-MLP Robustly Phyloclassifies Plastid Genomes to Late-Branching Cyanobacterial Clades

Using CYANO-MLP, we phyloclassified 437/440 (99.32%) plastid genomes to late-branching clades of Cyanobacteria, with 433 plastid genomes classifying to the B2+3 clade and four plastid genomes classifying to the A clade with high probabilities (Fig. 2; Table S5). Plastid genomes from all three Archaeplastida lineages phyloclassified to the late-branching cyanobacterial B2+3 clade. The majority of plastid bootstrap replicates classified to late-branching clades A, B1, and B2+3 with the median bootstrap frequency of all plastid groups against clade B2+3 at or above 70, except for the Glaucocystophyta genome (Fig. 2, S11-S14). Three remaining plastid genomes classified to early-diverging cyanobacterial clades; two to clade F and one to clade G (Fig 2, Table S5). With CYANO-MLP-BAL, 18 and 384 plastid genomes classified to clades A and B2+3 respectively (Tables S8).

Plastid genome classifications were mostly robust to model specifications. Classifications of clade-represented genomes were mostly unchanged in the CYANO-MLP[!A]) and CYANO-MLP[!C1] leave-clade-out models (Fig S21; Table S10,S11). Distinctly, plastid classifications with CYANO-MLP[!B1] were ambiguous with equal probabilities between clades A and B2+3 (Table S12). However, after retraining CYANO-MLP[!B1] using balanced training data (CYANO-MLP-BAL[!B]) phyloclassifications were restored to be similar to those with CYANO-MLP and CYANO-MLP-BAL (Table S12,S13). We then developed two phyloclassifers including training data only from late-branching clades A, B1, and B2+3, one with balanced training data (CYANO-MLP-BAL[AB1B2+3]) and one without (CYANO-MLP[AB1B2+3]). CYANO-MLP[AB1B2+3] classified plastids equivocally between clades A and B2+3, similarly to CYANO-MLP[!B1], though slightly favoring clade B2+3 (Table S13,S14). After balancing data by resampling, CYANO-MLP-BAL[AB1B2+3] more decisively phyloclassified 331 plastid genomes to clade B2+3 and only 106 genomes to clade A (Table S8,S13,S14). Remarkably, phyloclassifications of both plastid and B2+3-cyanobacterial genomes with the CYANO-MLP[!B2+3] leave-clade-out model were equivocal and similar to one another (Fig S21; Table S6-S9), consistent with “none-of-the-above” classification and providing compelling additional support that plastids belong to clade B2+3.

### Phyloclassification of *G. lithophora* is Consistent with its Early Divergence within Cyanobacteria

Recent phylogenomic analyses support a sister relationship between plastids and an early-diverging lineage containing *G. lithophora* as its only member [43, 52]. With only one genome, there was insufficient tRNA sequence data to estimate CIFs for this lineage. Instead, we classified the *G. lithophora* genome using CYANO-MLP to determine if it classified similarly to plastids, which would be consistent with a sister relationship of *G. lithophora* and plastids. We found that the *G. lithophora* genome obtained greater than 75% total classification probability against three early-diverging clades, classifying to clade F with probability 57.3%, to clade G with probability 18.4%, and to clade E with probability 3.2%. In addition, *G. lithophora* classified to the late-diverging clade A with probability 20.3% (Fig. 2). We interpreted the results as consistent with a “none-of-the-above” classification, yet, favoring an early-branching of *G. lithophora*, in agreement with recent phylogenomic analyses [43, 52]. Notably, the incongruity of our results for *G. lithophora* and plastids rejects their sister relationship.

### Inadequate modeling of systematic biases can explain discrepancies with prior work

We examined goodness of fit of various evolutionary models to published combined cyanobacterial/plastid phylogenomic datasets by posterior predictive analysis [10, 28]. We found evidence that site-specific amino acid constraints are critical to fitting all three cyanobacterial/plastid phylogenomic datasets (Figure 3A; Table S15). The empirical matrix model LG+4Γ [33], with site-rate heterogeneity, fails to model site-specific substitution processes [28, 29] in all three phylogenomic datasets and fits them poorly (Fig. 3A, Table S15). The inadequacy of empirical matrix models to fit data with site-specific constraints was previously reported [29]; their use to fit such data results in long-branch attraction artifacts caused by underestimation of homoplasy [28]. In contrast, the CAT model [28, 29] specifically accommodates site-specific constraints, fitting all three datasets adequately (Fig. 3A; Table S15). However, even in combination with CAT, none of the amino acid recoding methods adequately mitigate lineage-specific compositional biases (*Z*≥5, Fig. 3B; Table S15). When lineage-specific compositional biases are not adequately modeled, unrelated sequences with similar compositions may artifactually cluster during phylogenetic tree reconstruction [10].

**Figure 3.**
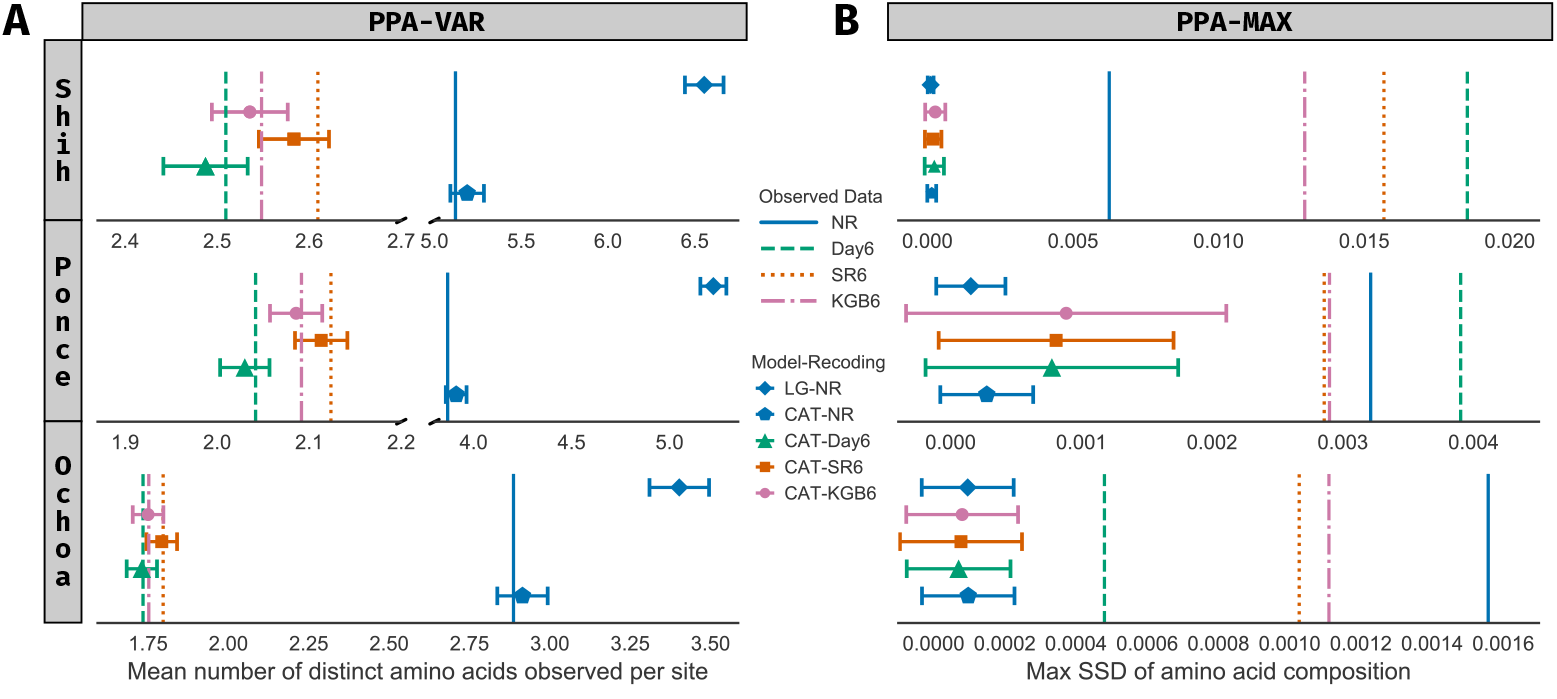
Results of posterior predictive analyses of phylogenomic datasets of Shih [47], Ponce-Toledo [43], and Ochoa de Alda [39]. Rows within each panel correspond to phylogenomic datasets. Observed values calculated for each test statistic are represented by vertical lines. Color and patterns of vertical lines indicate amino acid recoding strategies, respectively NR: No recoding, DAY6: six-state Dayhoff recoding, SR6: the six-state recoding strategy of [51], and KGB6: the six-state recoding strategy of [27]. Symbols show average values for two posterior predictive test statistics calculated from simulated datasets, with error bars showing ± five standard deviations. Symbol shapes correspond to phylogenetic models (LG: LG+4Γ, CAT: CAT-GTR+4Γ) and symbol colors show recoding strategies for simulated data. If similarly colored error bars contain vertical lines, the given phylogenetic model adequately describes systematic biases of the given datas. (A) Results with the PPA-VAR statistic [28] assessing fit of models to site-specific constraints in the data. (B) Results with the PPA-MAX statistic, measured as squared standard deviation (SSD) of amino acid composition [10], showing generally poor fit of models against the lineage-specific compositional biases in the data.

## Discussion

We recovered strong support for a late-branching origin of plastids within or closely related to the B2+3 clade of the CyanoToL (Figs 1-2; Table S5). Furthermore, our result of a late-branching clade B2+3 origin of plastids is robust to bootstrap resampling of tRNA structural positions (Fig 2; Supplemental file 1), missing data (Fig S15-S18,S21; Table S7-S12), and unbalanced training data (Fig S19-S21; Table S7,S8,S12,S13). Additionally, we were able to reject recent hypotheses supporting the early-branching *G. lithophora* as sister to plastids [43, 52] (Fig. 2). Our results conform to independent metabolic evidence that plastids originated from a unicellular starch-producing diazotrophic cyanobacterial species [6, 15], and independent comparative evidence that photosynthetic eukaryotes originated and diversified rapidly in a low-salinity habitat [8, 52].

Importantly, the significantly lower classification accuracy of CYANO-MLP on class-permuted training datasets (Fig. S9) support that CYANO-MLP phyloclassifications depend on learned phylogenetic signals in cyanobacterial tRNA CIFs. Furthermore, we argue against the interpretation that plastid genomes have experienced distinctive selection pressures yielding idiosyncratic score vectors and artifactual results, because of the consistency with which plastids classified in the various re-specifications of CYANO-MLP, and consistent classifications of C1 clade Cyanobacteria with reduced genomes and the *P. chromatophora* chromatophore genome, presumably under similar selection pressures as plastid genomes, to the C1 clade (Fig. in concert with previous work [39, 47, 52].

To reconcile recent studies with our results, we reexamined the fits of recently used models and recoding strategies to three published cyanobacterial/plastid phylogenomic datasets (Fig. 3; Table S15). We found that the CAT model [28, 29] accommodated site-specific constraints (Fig. 3A; Table S15), however, amino acid recoding strategies were unable to mitigate lineage-specific compositional biases (Fig. 3B; Table S15). Only one prior phylogenomic study took into account both sources of bias [39], in which 16S rDNA nucleotide data was modeled using CAT-GTR while removing compositionally divergent taxa to achieve compositional homogeneity. Notably, the findings of [39] are consistent with ours in supporting a late-branching origin of plastids within Cyanobacteria.

## Supporting Information

**Figure S1.**
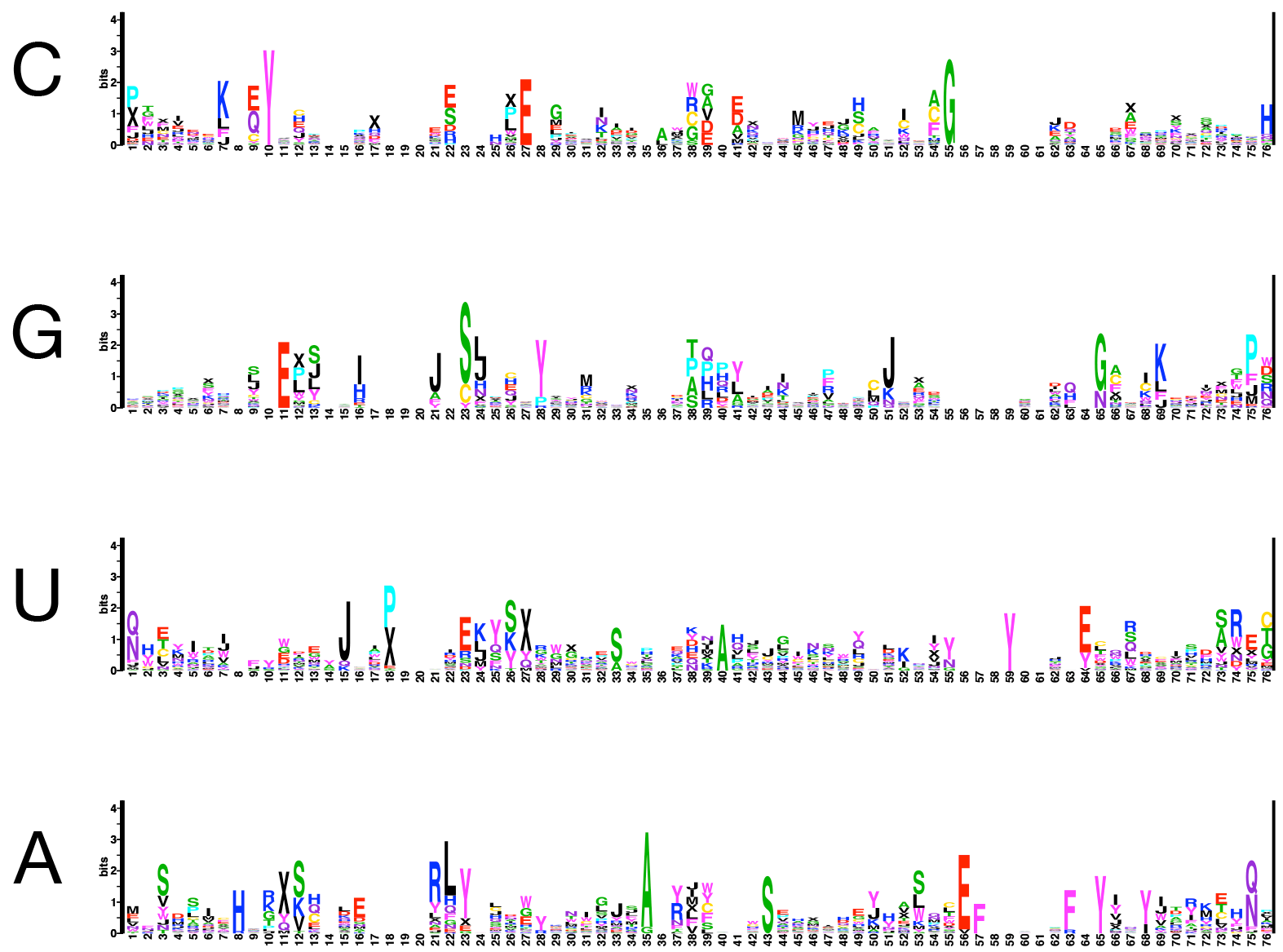
Function Logos for Cyanobacterial Clade A

**Figure S2.**
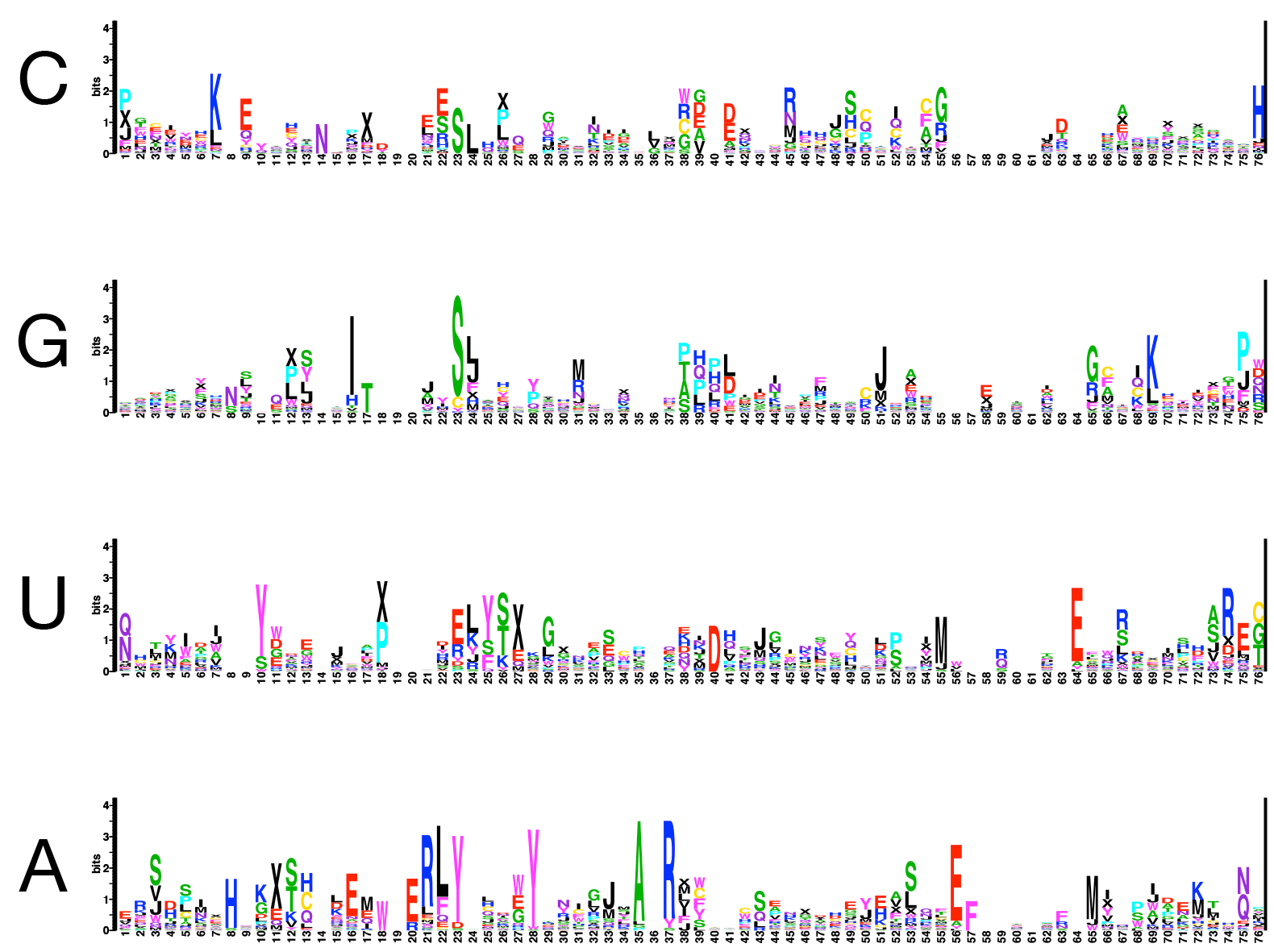
Function Logos for Cyanobacterial Clade B1

**Figure S3.**
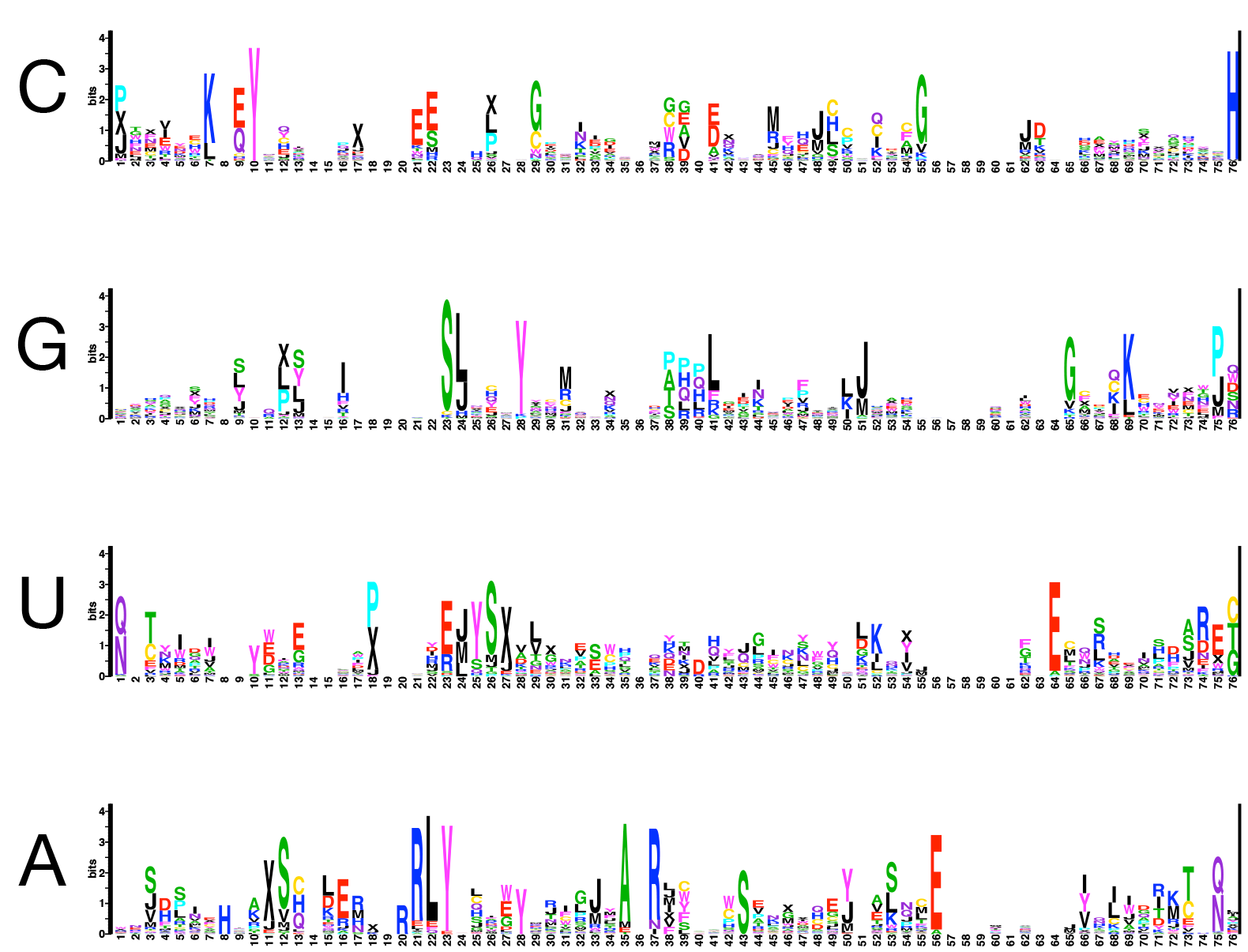
Function Logos for Cyanobacterial Clade B2+3

**Figure S4.**
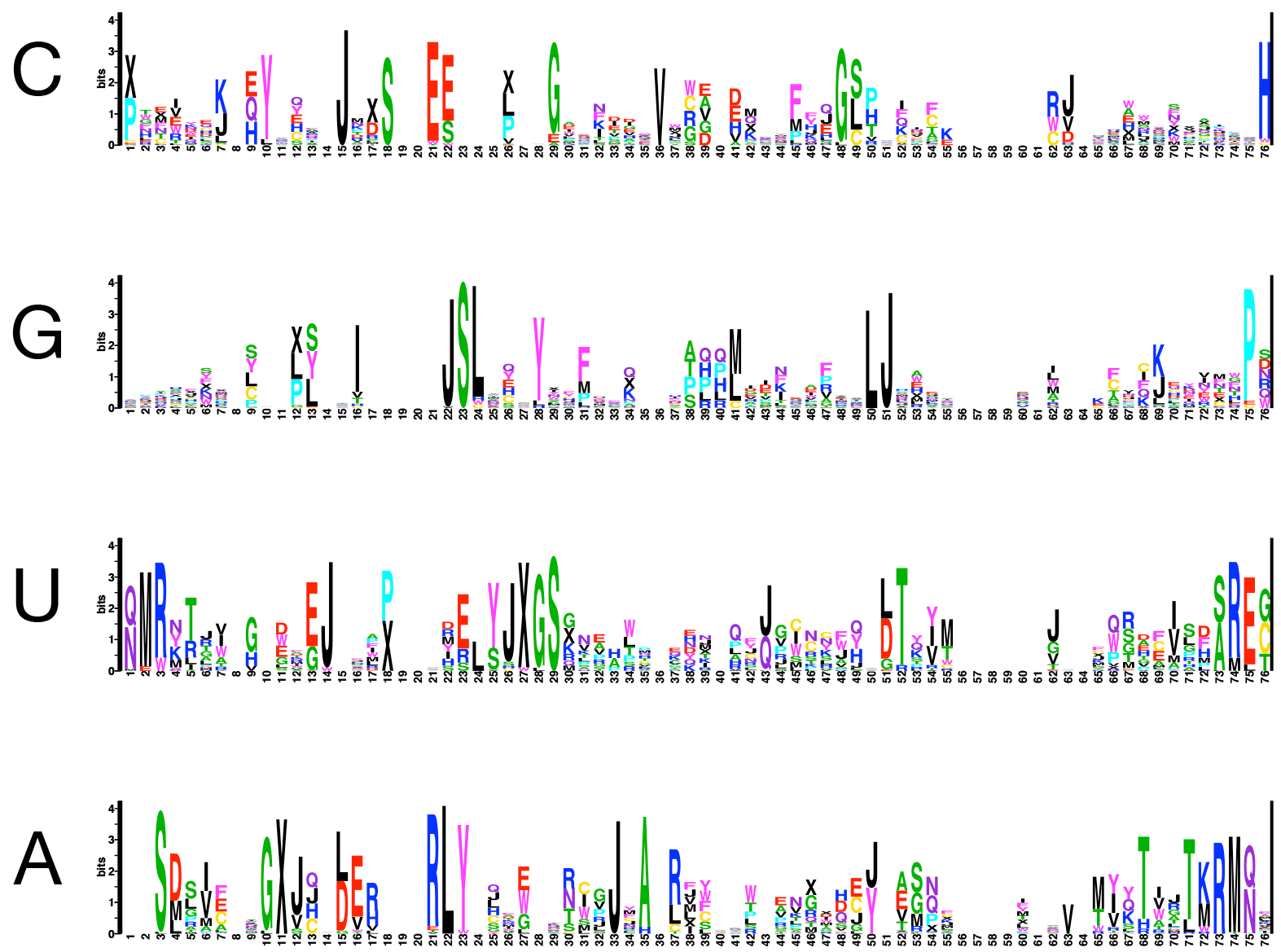
Function Logos for Cyanobacterial Clade C1

**Figure S5.**
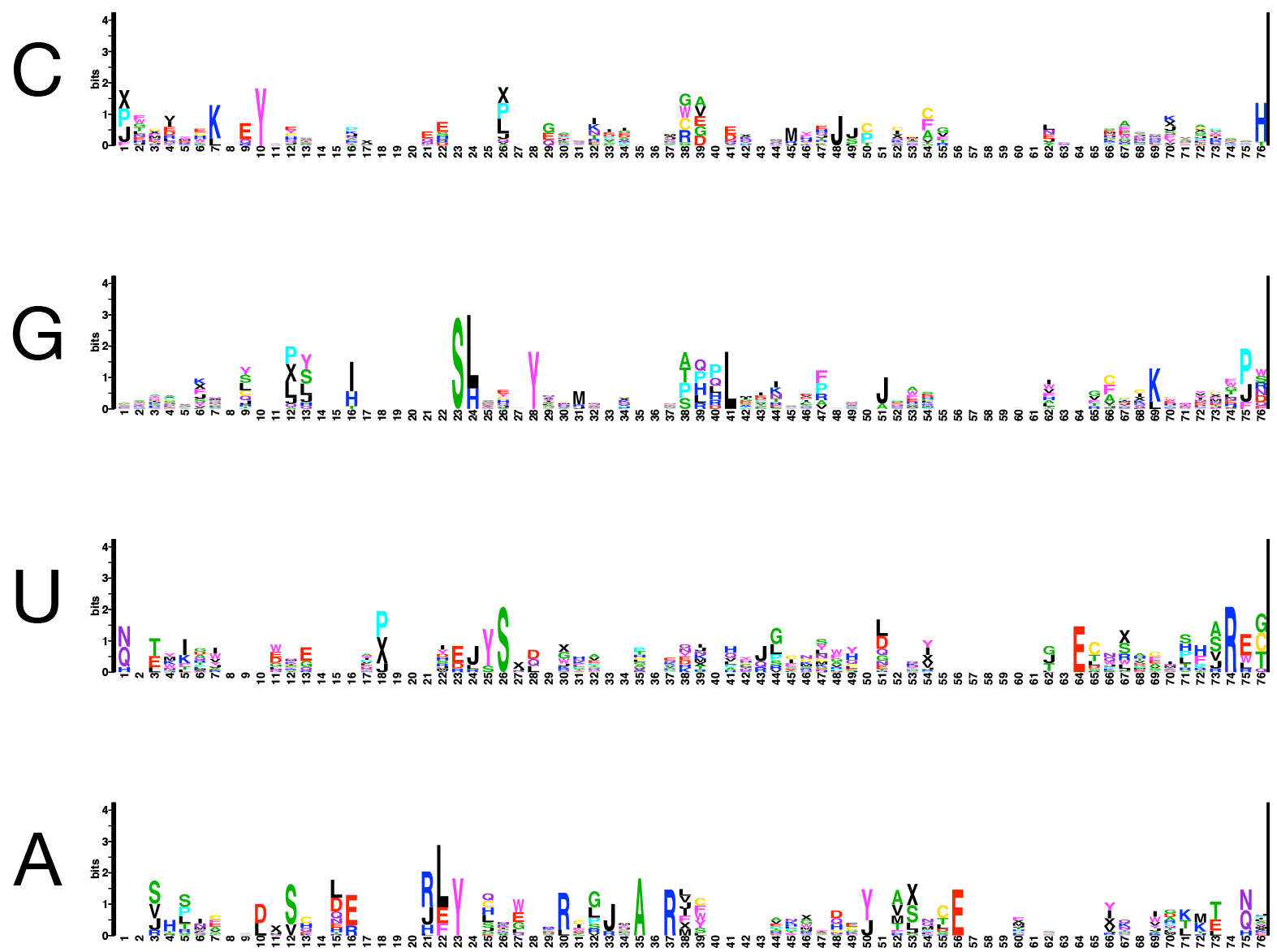
Function Logos for Cyanobacterial Clade C3

**Figure S6.**
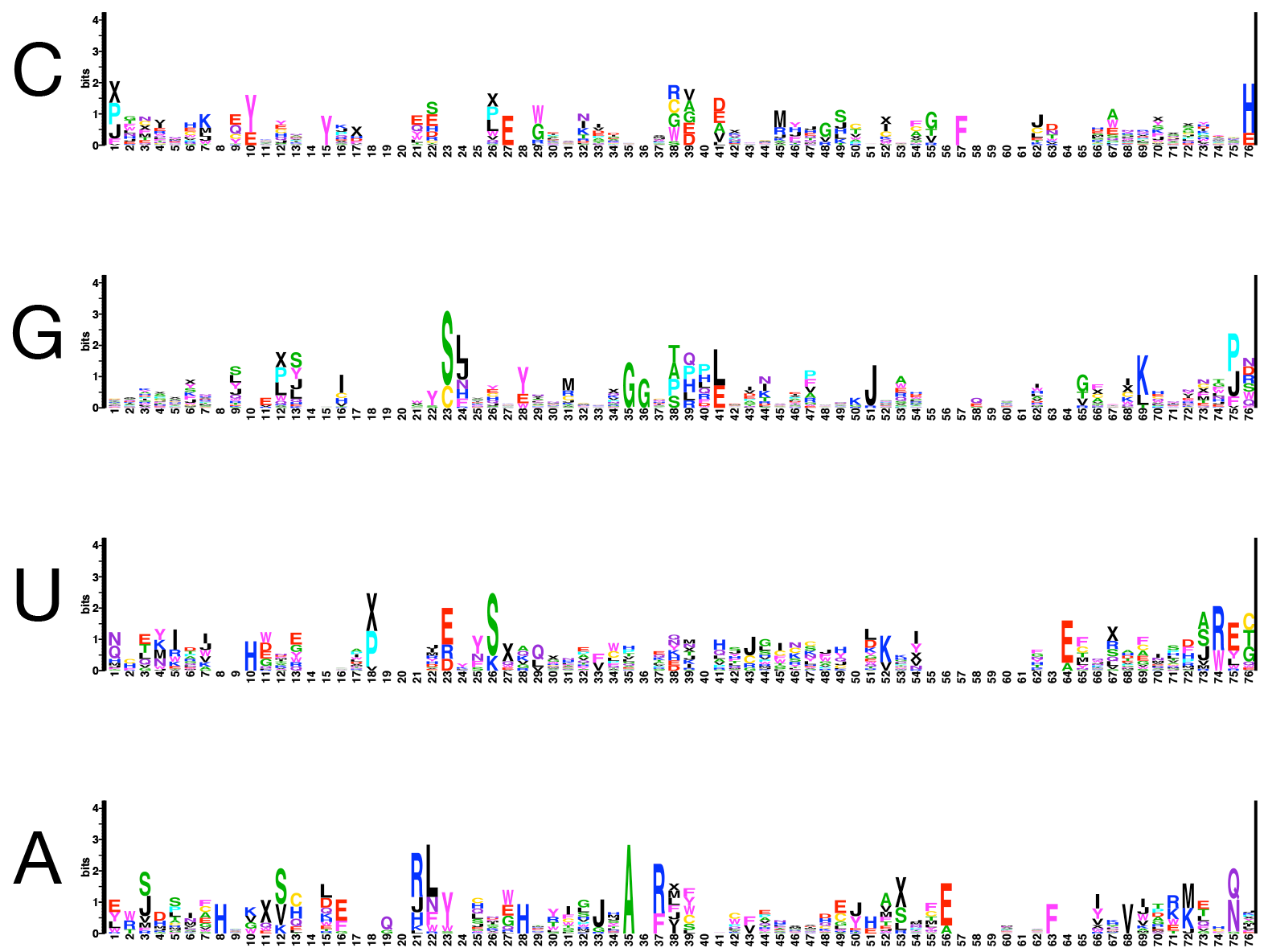
Function Logos for Cyanobacterial Clade E

**Figure S7.**
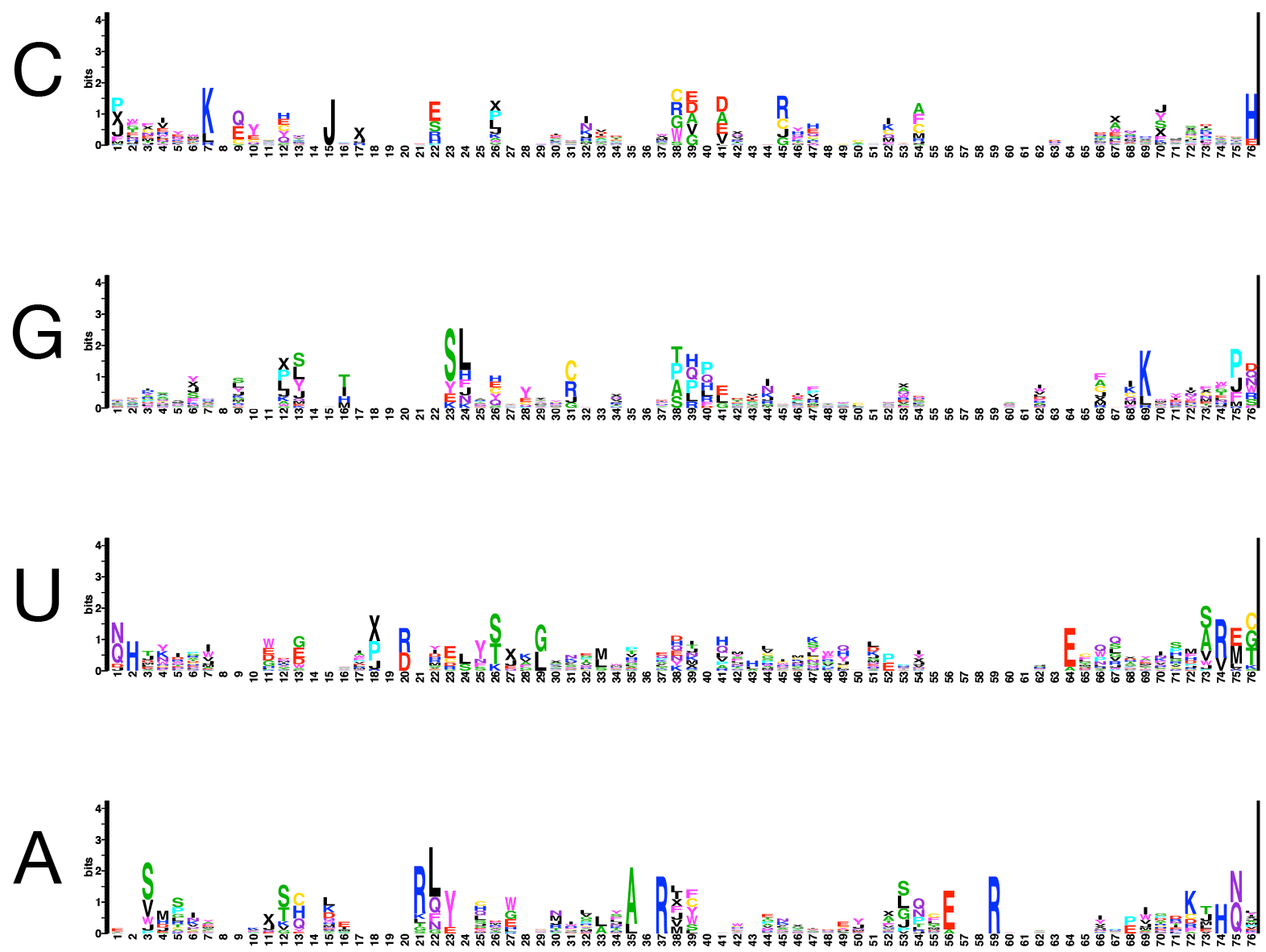
Function Logos for Cyanobacterial Clade F

**Figure S8.**
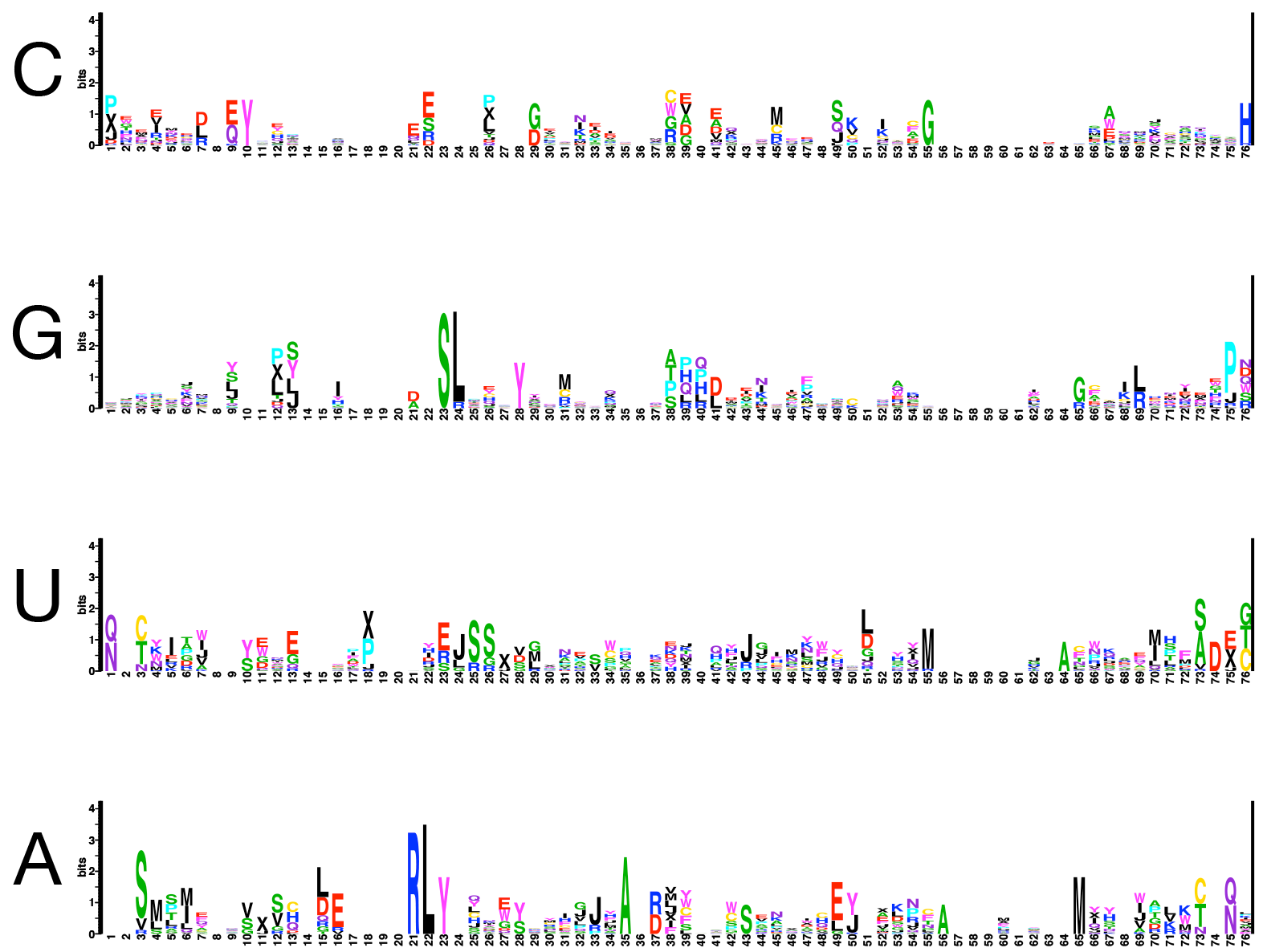
Function Logos for Cyanobacterial Clade G

**Table S1.**
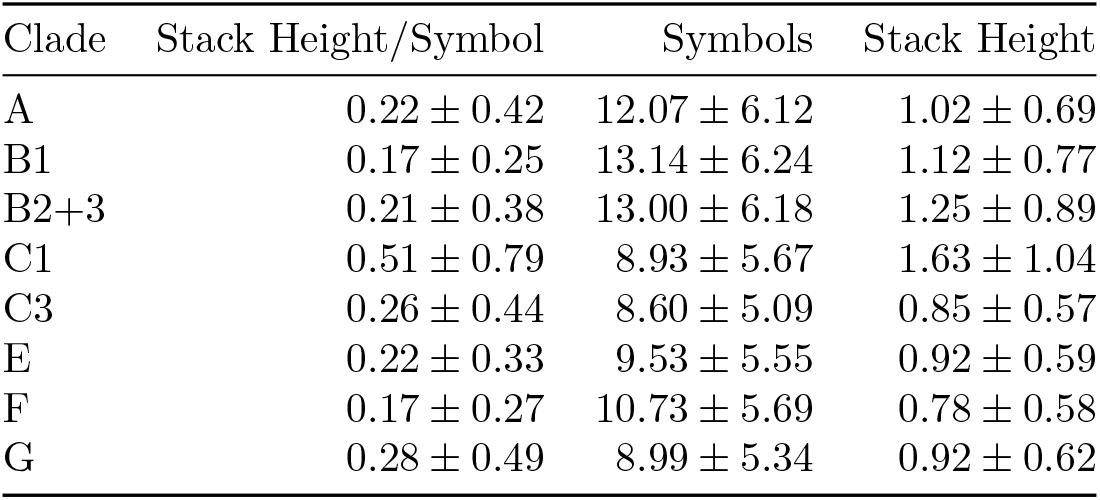
Descriptive statistics of Cyanobacterial clade function logos. (Stack Height/Symbol) Average of information content for each site divided by the number of symbols for that site, (Symbols) Average number of symbols per site, and (Stack Height) the average information content in bits of each site. Sites with zero information were excluded from calculations.

**Table 2.**
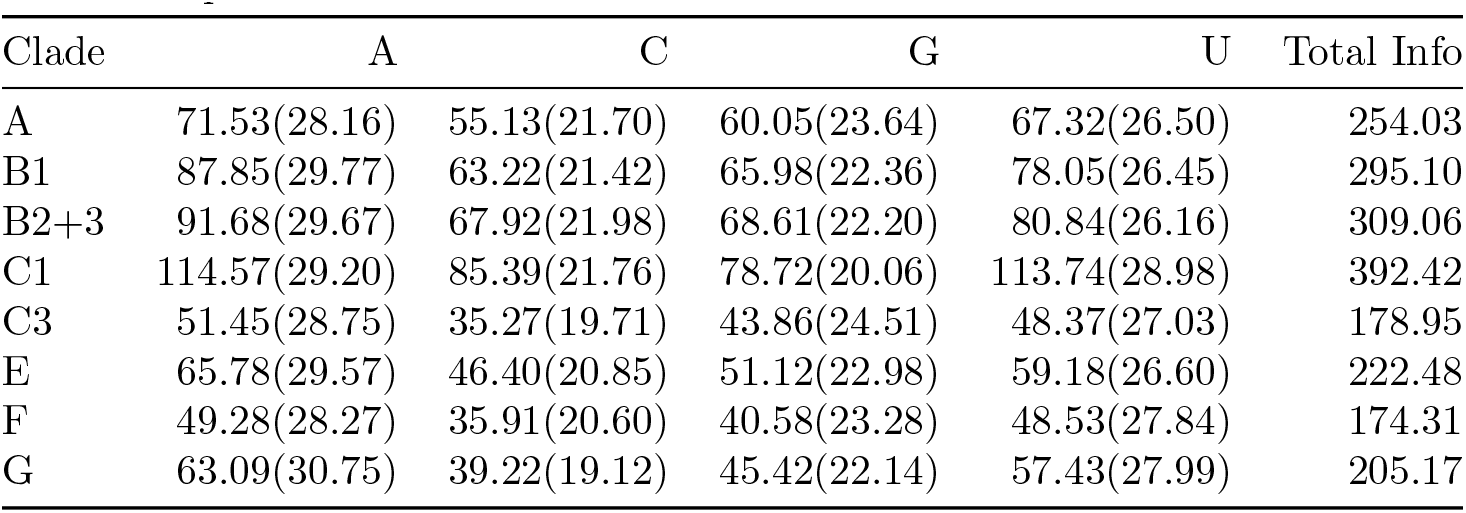
Information content per nucleotide for each Cyanobacterial clade. Number in parenthesis is percent of total information.

**Figure S9.**
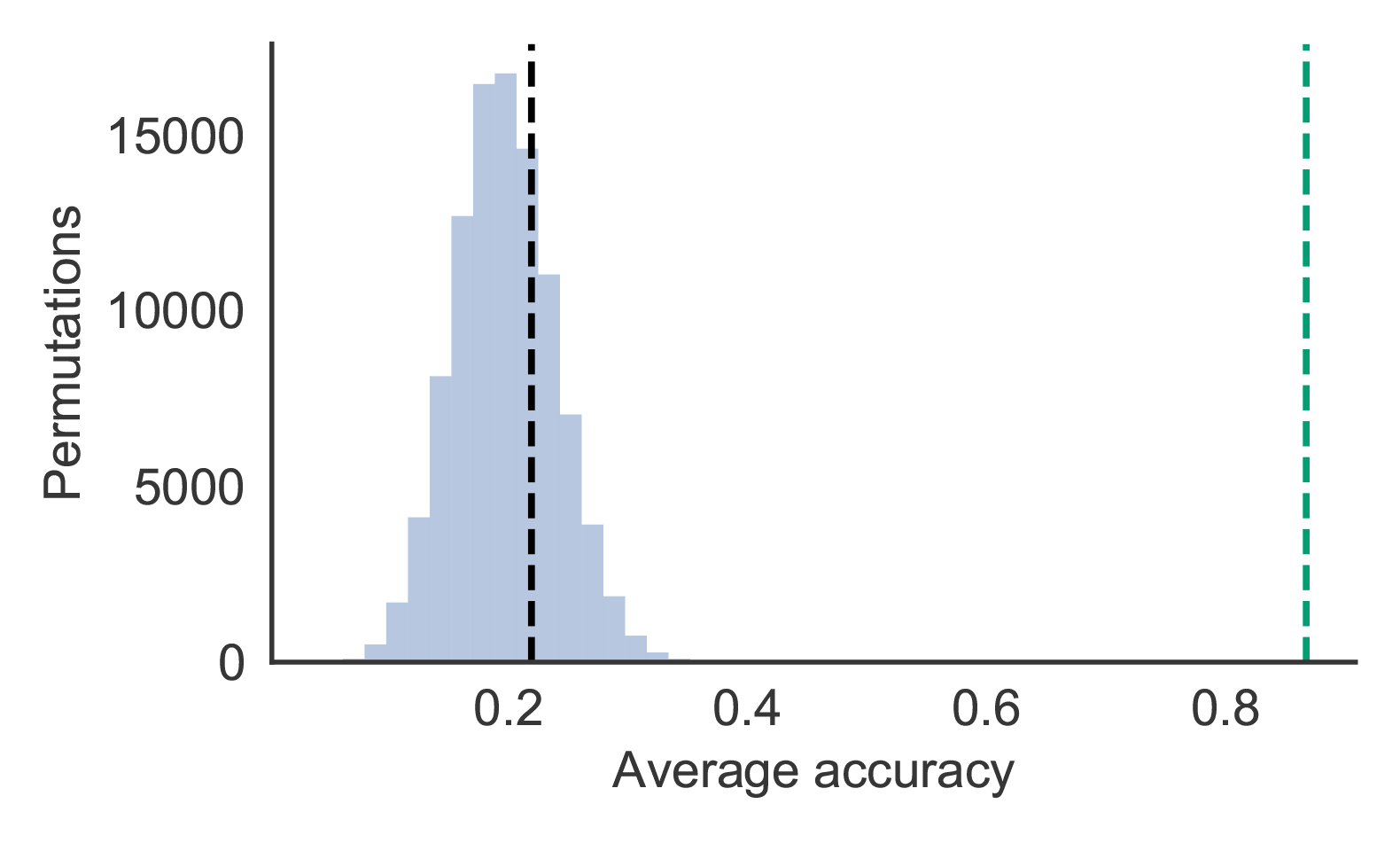
Null distribution of average accuracy using LOOCV estimated by 100,000 label swapping permutation datasets. Black dotted line is the expected average accuracy if cyanobacteria genomes were randomly classified. Green dotted line is the average accuracy using our single hidden layer phyloclassifier.

**Figure S10.**
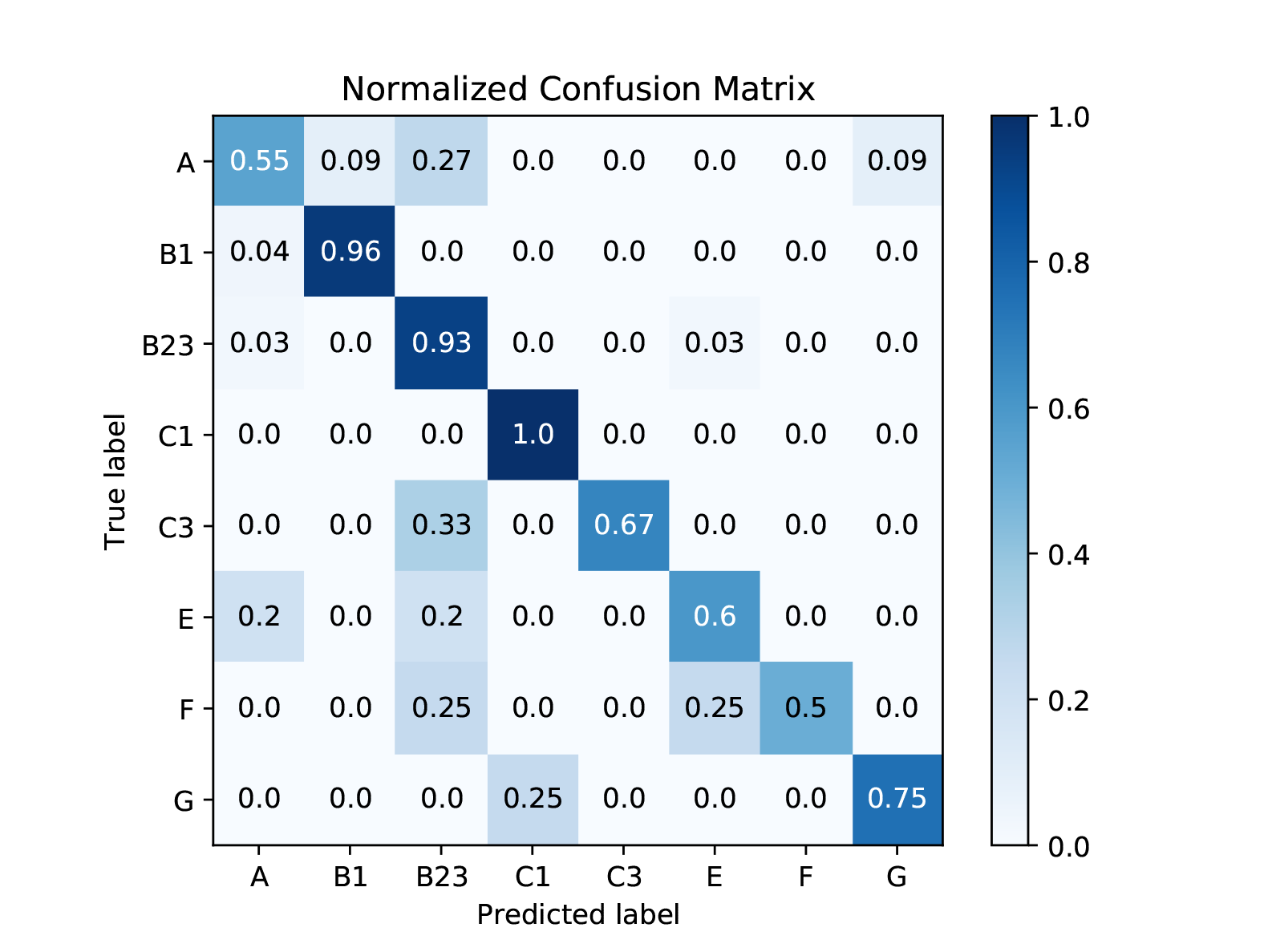
Normalized Confusion Matrix for CYANO-MLP

**Figure S11.**
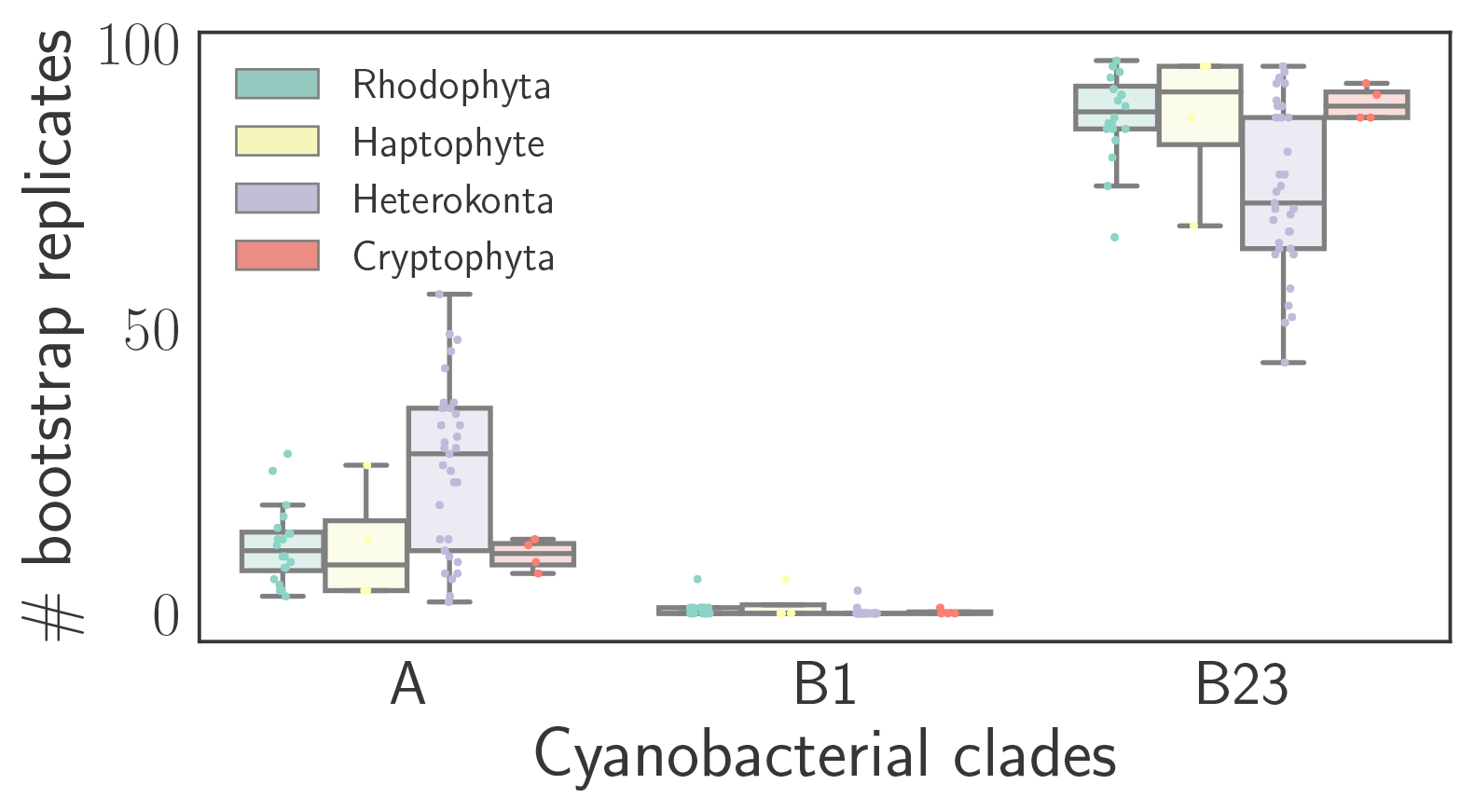
Classification results of 100 bootstrap replicates of each Rhodophyta derived plastid genome. Results are summarized by Red plastid group with boxes spanning from the 25th percentile (bottom) to the 75th percentile (top) of bootstrap replicates classifying to the indicated Cyanobacterial clade per genome with the bisecting line marking the median value. Error bars indicate the shorter of either ± the interquartile range or the span of bootstrap replicates per genome. Dots show bootstrap replicates for individual genomes. Cyanobacterial clades C1, C3, E, F, and G were omitted because a limited number of bootstrap replicates per genome classified to these clades (see Fig. S13).

**Figure S12.**
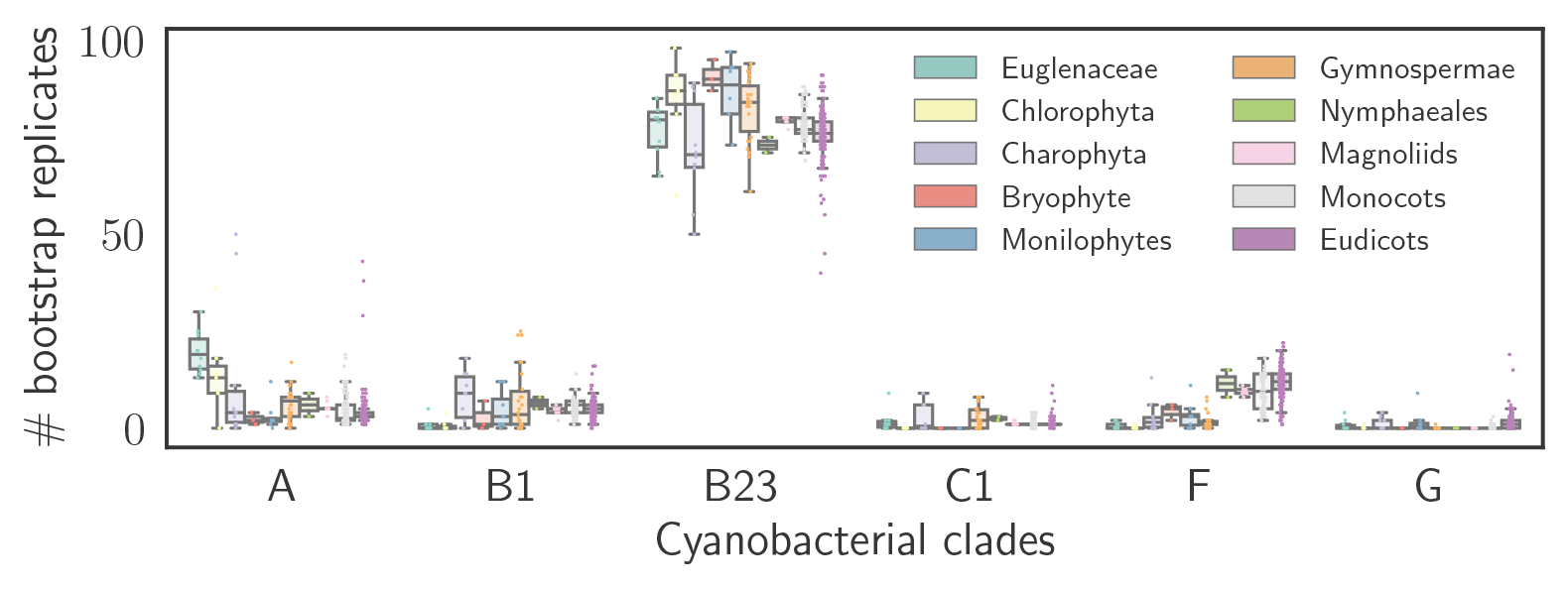
Classification results of 100 bootstrap replicates of each Chloroplastida derived plastid genome. Cyanobacterial clades C3 and E were omitted because a limited number of bootstrap replicates per genome classified to these clades (see Fig. S14). Results are summarized identically to Fig. S11.

**Figure S13.**
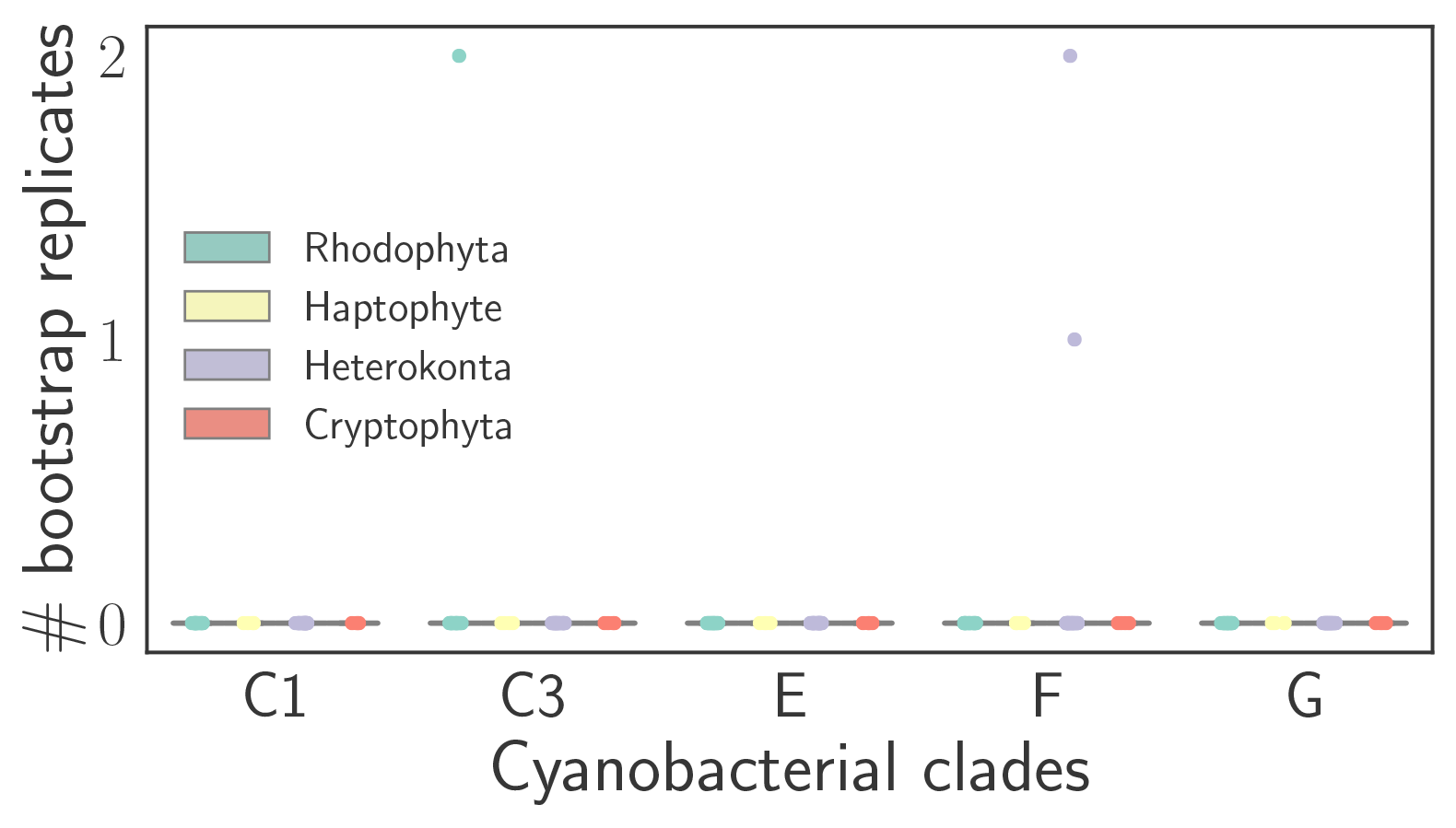
Classification results of 100 bootstrap replicates of each RED plastid genome for cyanobacterial clades C1, C3, E, F, and G. Results are summarized identically to Figure S11.

**Figure S14.**
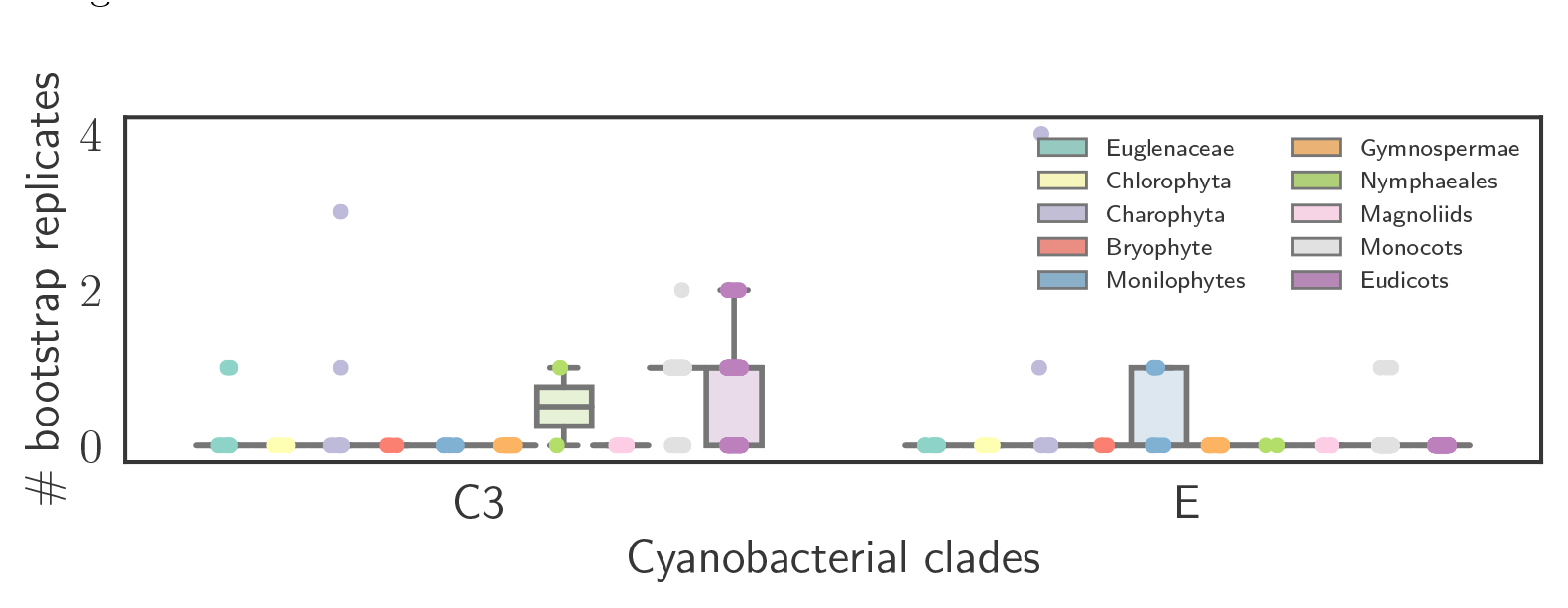
Classification results of 100 bootstrap replicates of each GREEN plastid genome for cyanobacterial clades C3 and E. Results are summarized identically to Figure S11.

**Figure S15.**
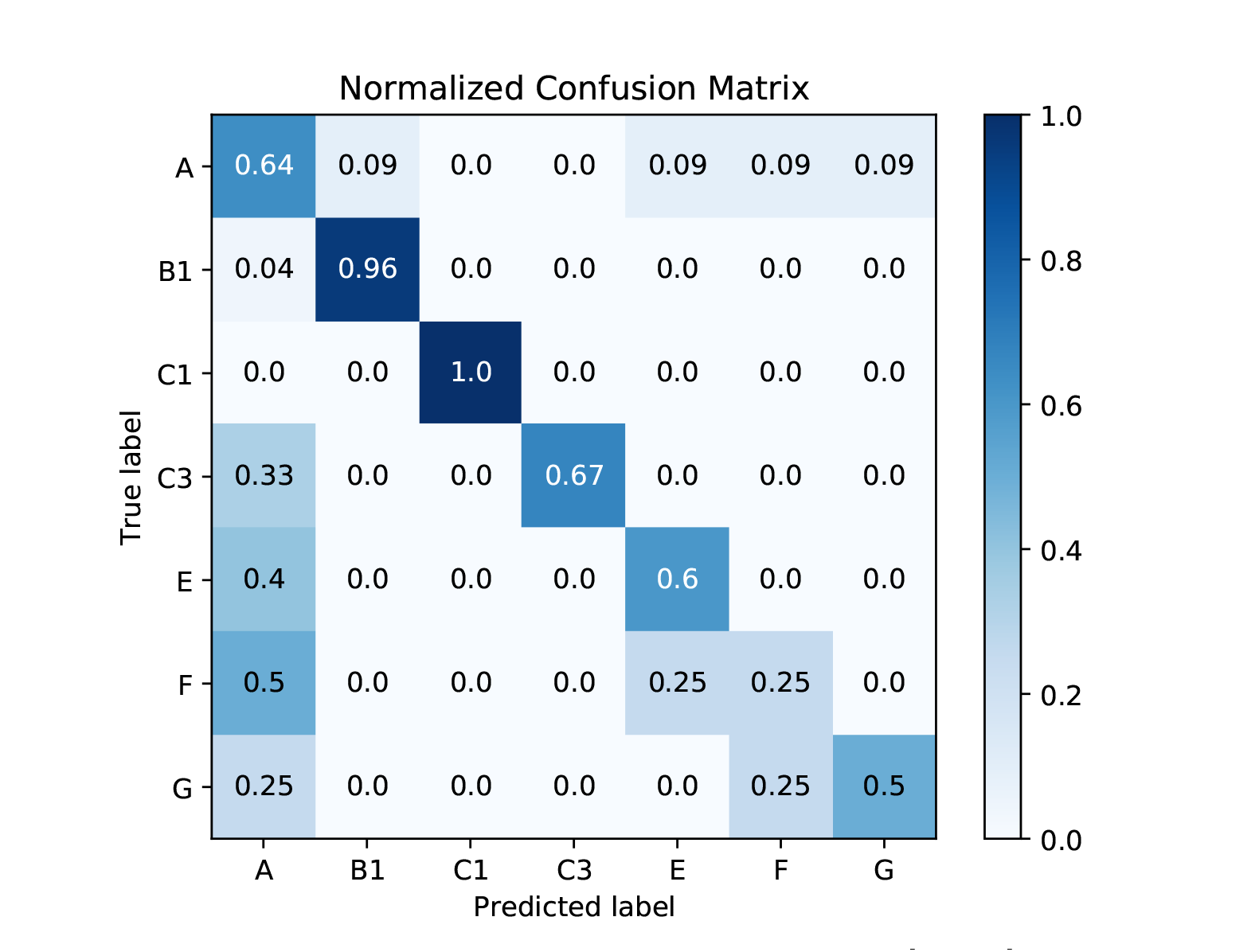
Normalized Confusion Matrix for CYANO-MLP[!B2+3]

**Figure S16.**
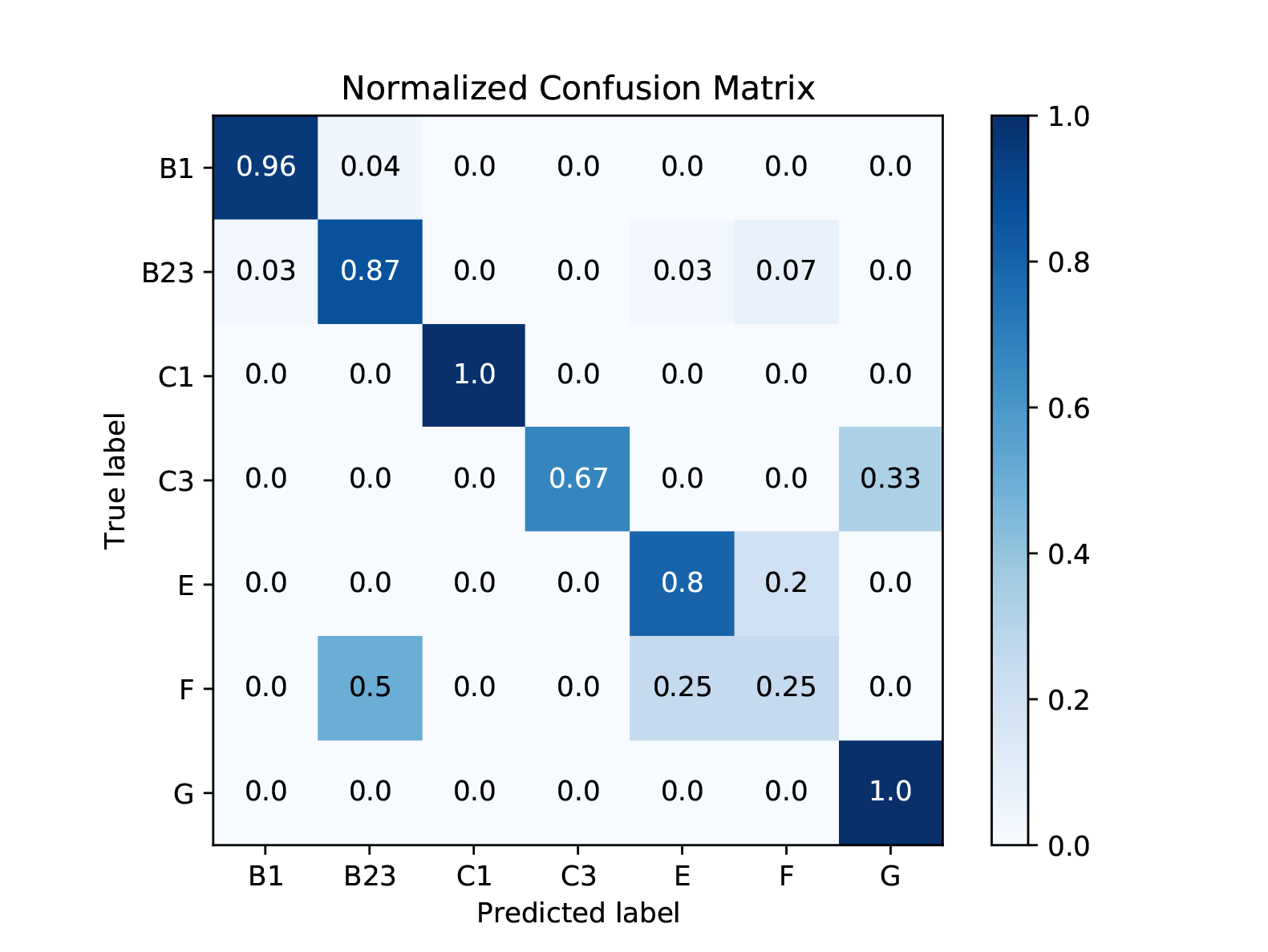
Normalized Confusion Matrix for CYANO-MLP[!A]

**Figure S17.**
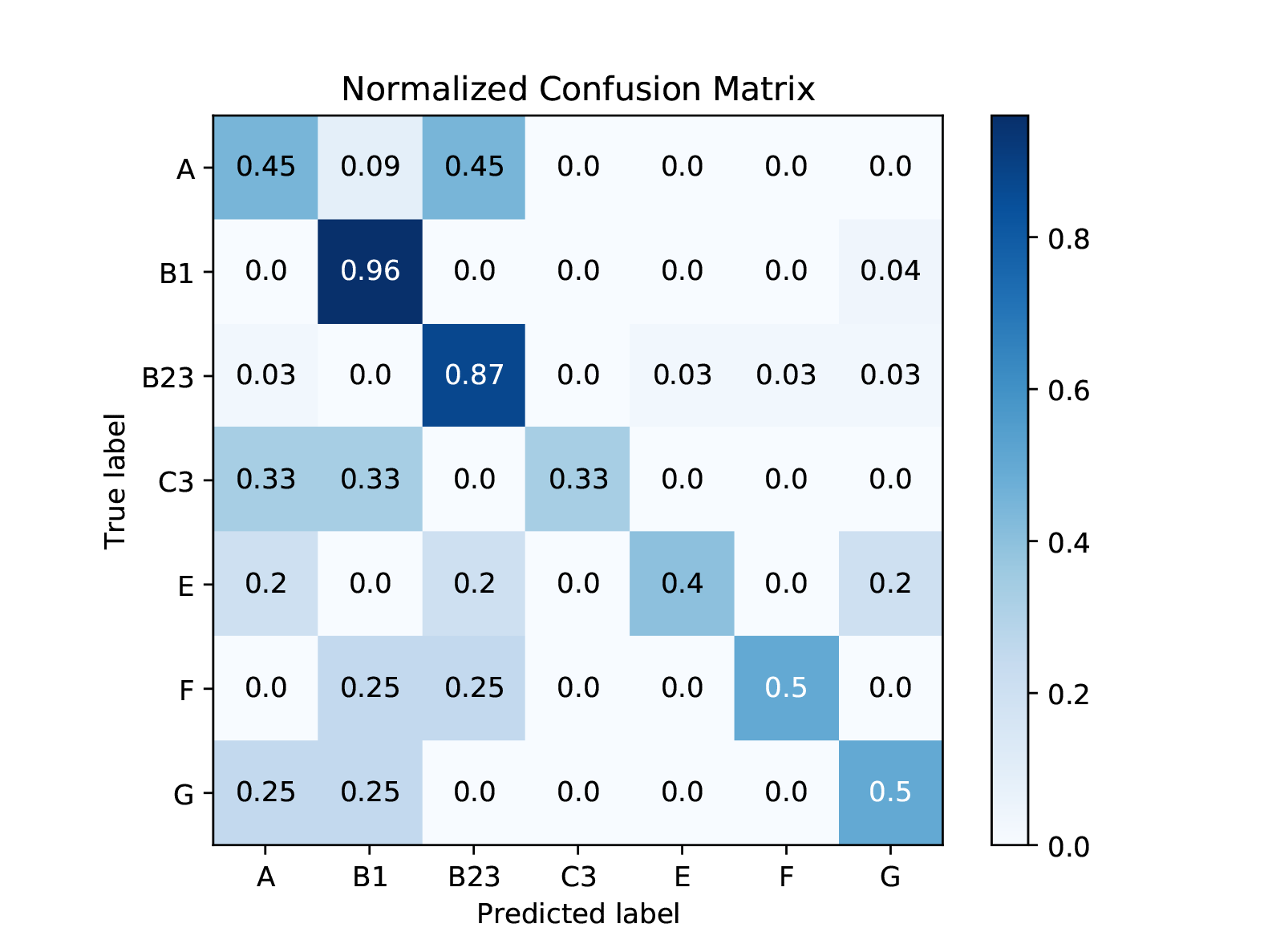
Normalized Confusion Matrix for CYANO-MLP[!C1]

**Figure S18.**
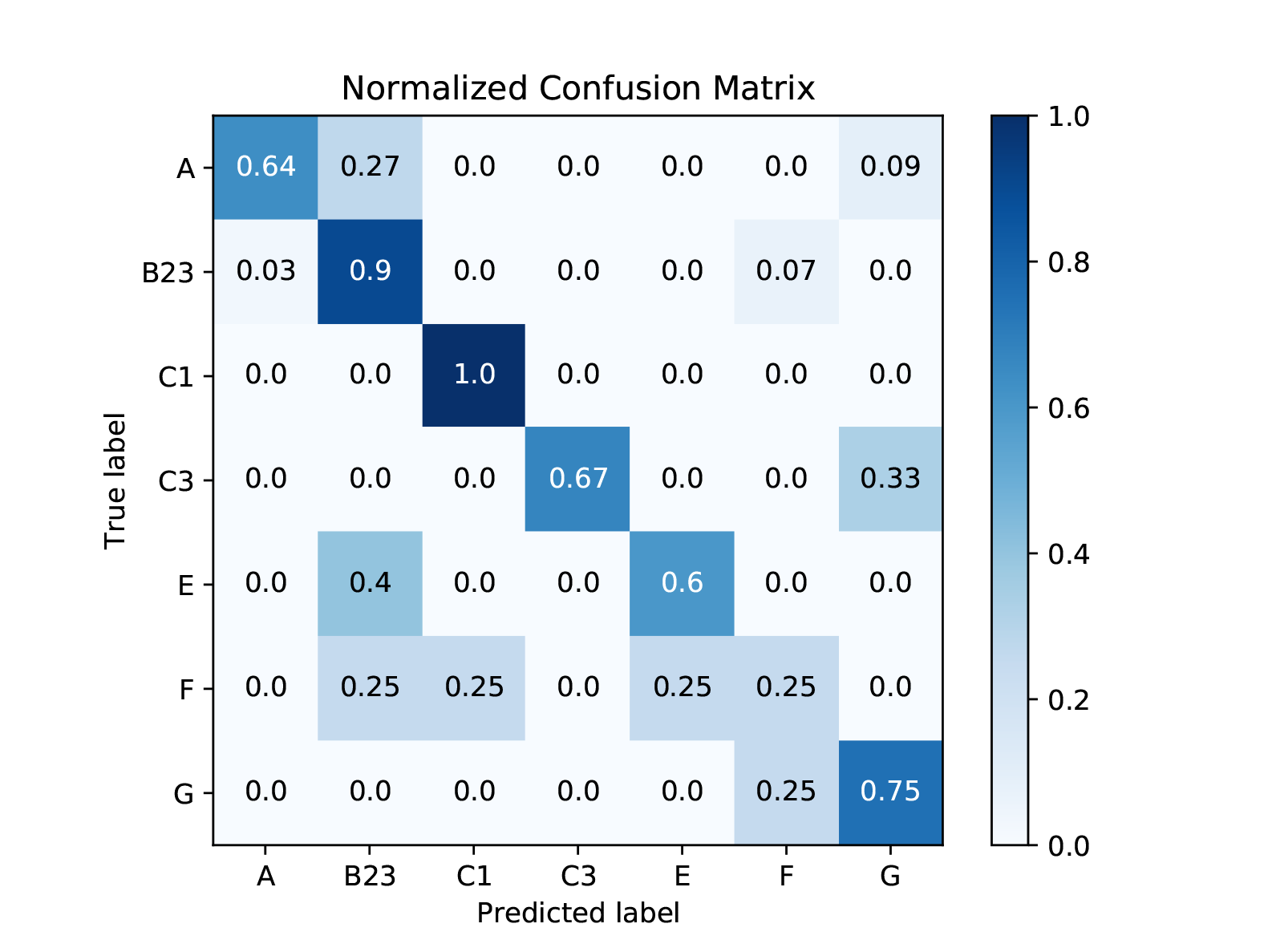
Normalized Confusion Matrix for CYANO-MLP[!B1]

**Figure S19.**
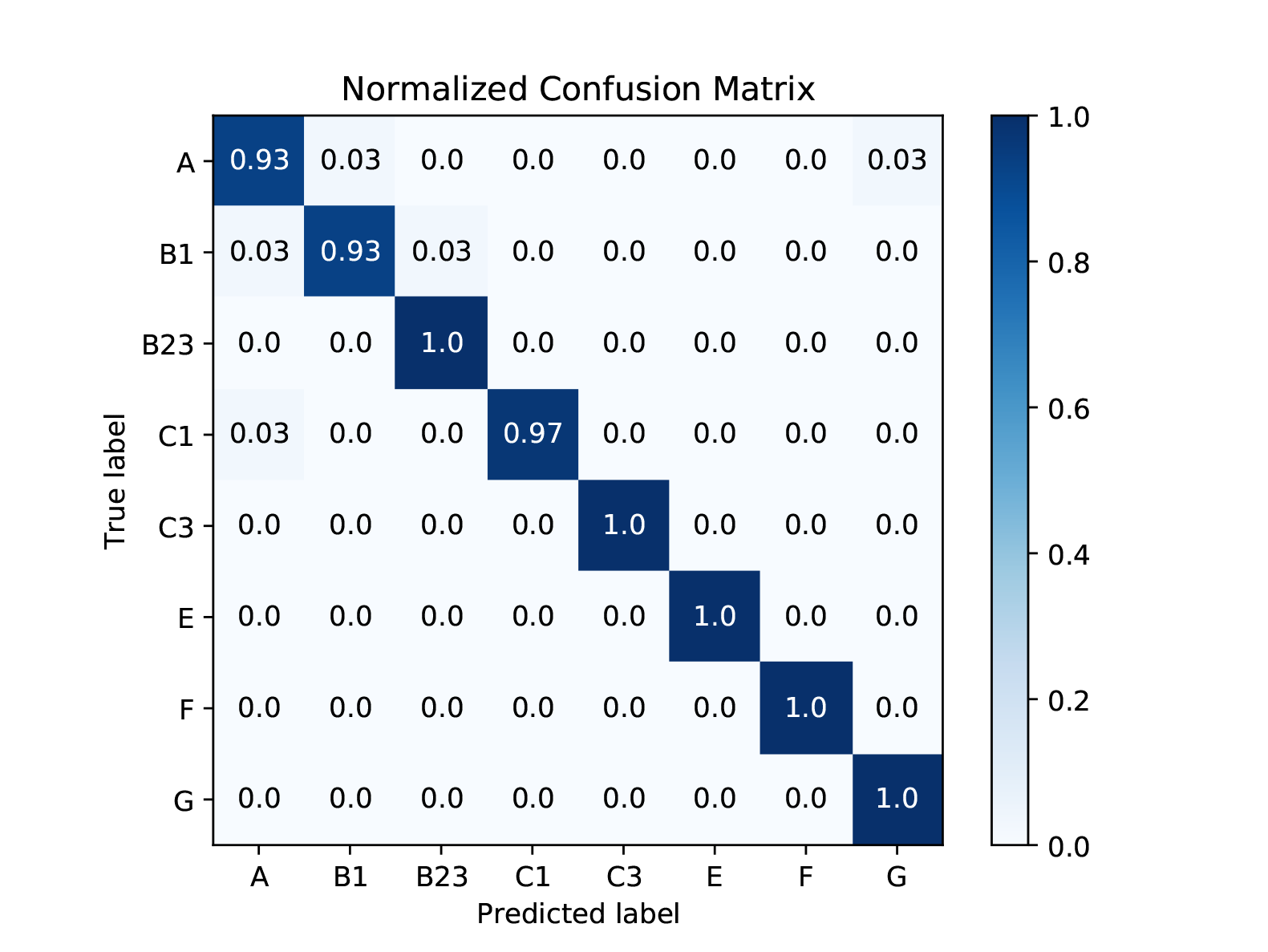
Normalized Confusion Matrix for CYANO-MLP-BAL

**Figure S20.**
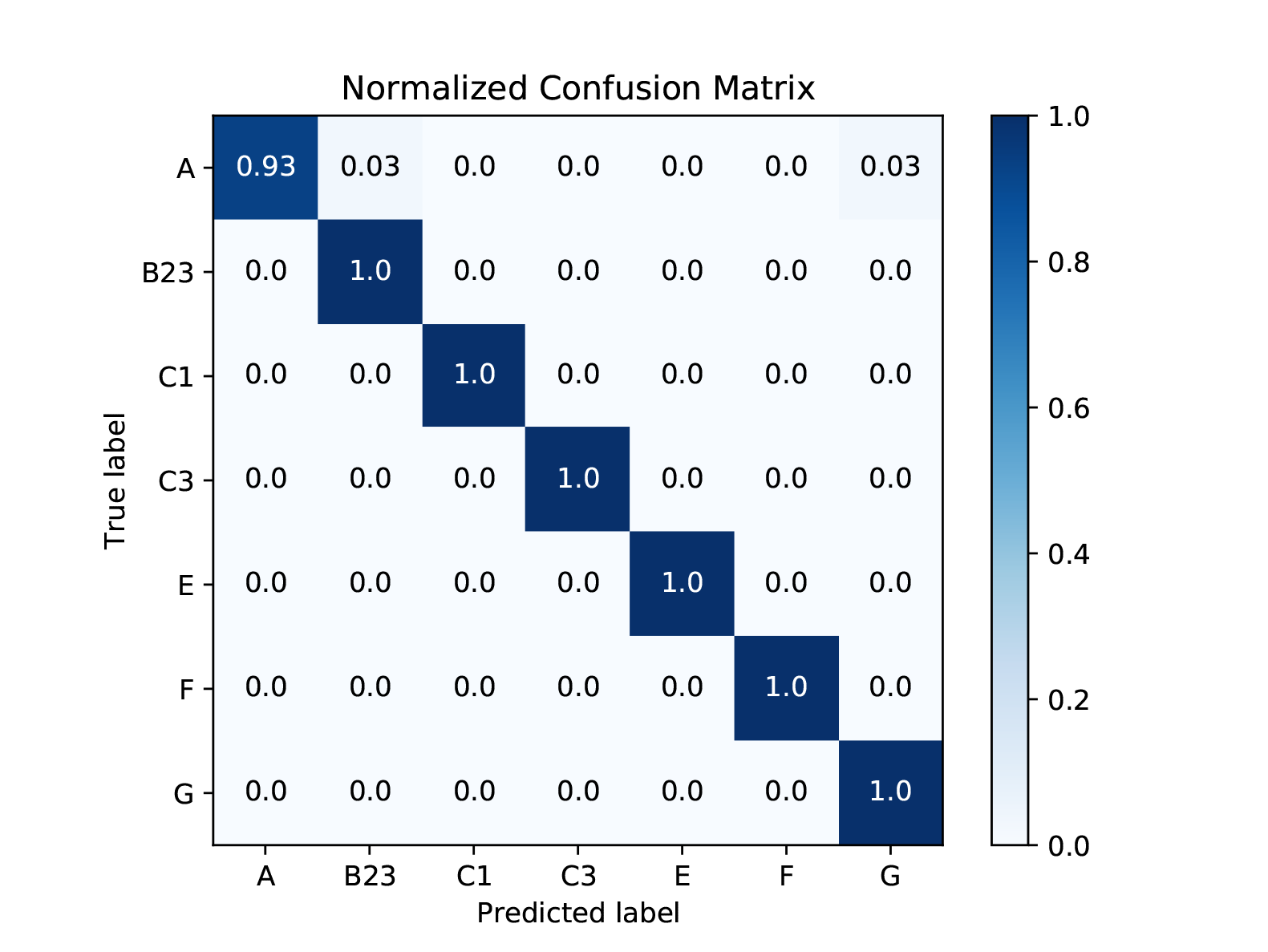
Normalized Confusion Matrix for CYANO-MLP-BAL[!B1]

**Table 3.**
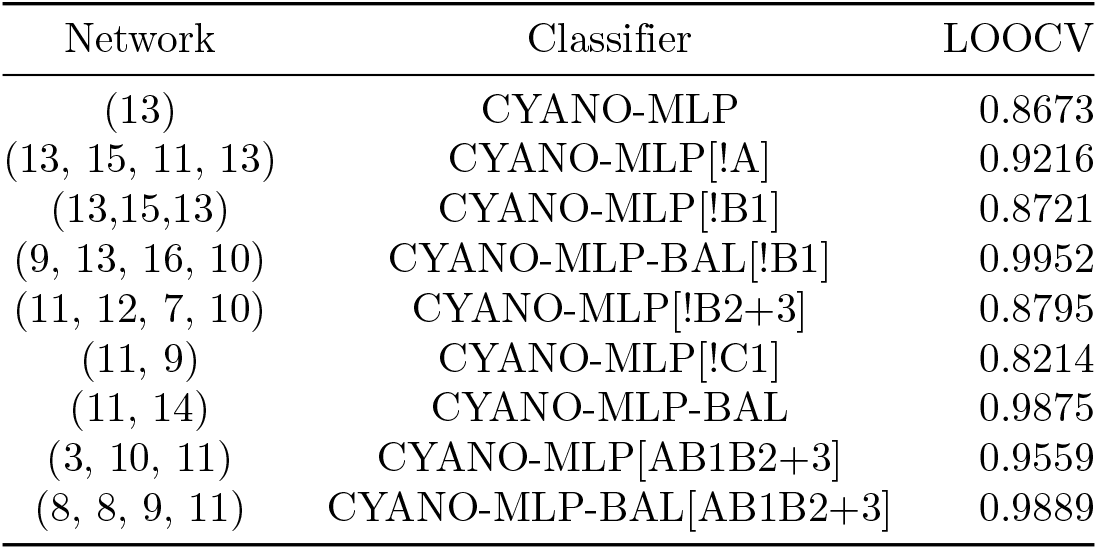
Network architecture and average accuracy using Leave-One-Out Cross-Validation for CYANO-MLP, .

**Table 4.**
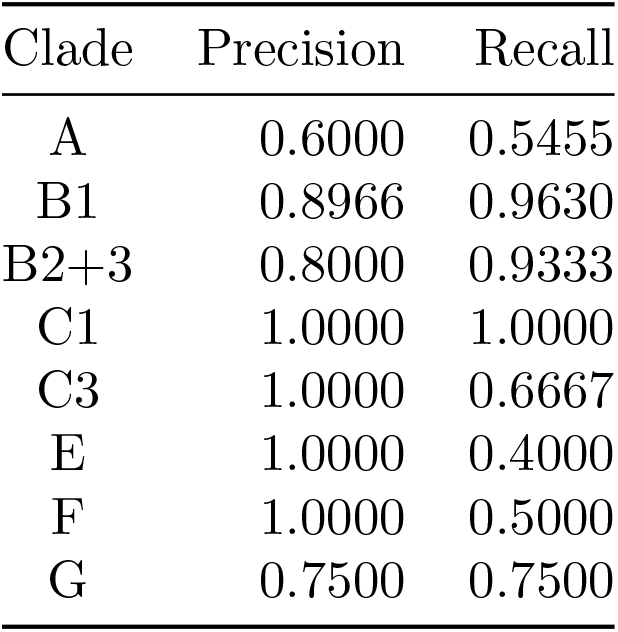
Precision and Recall for CYANO-MLP for each Cyanobacterial clade.

**Table 5.**
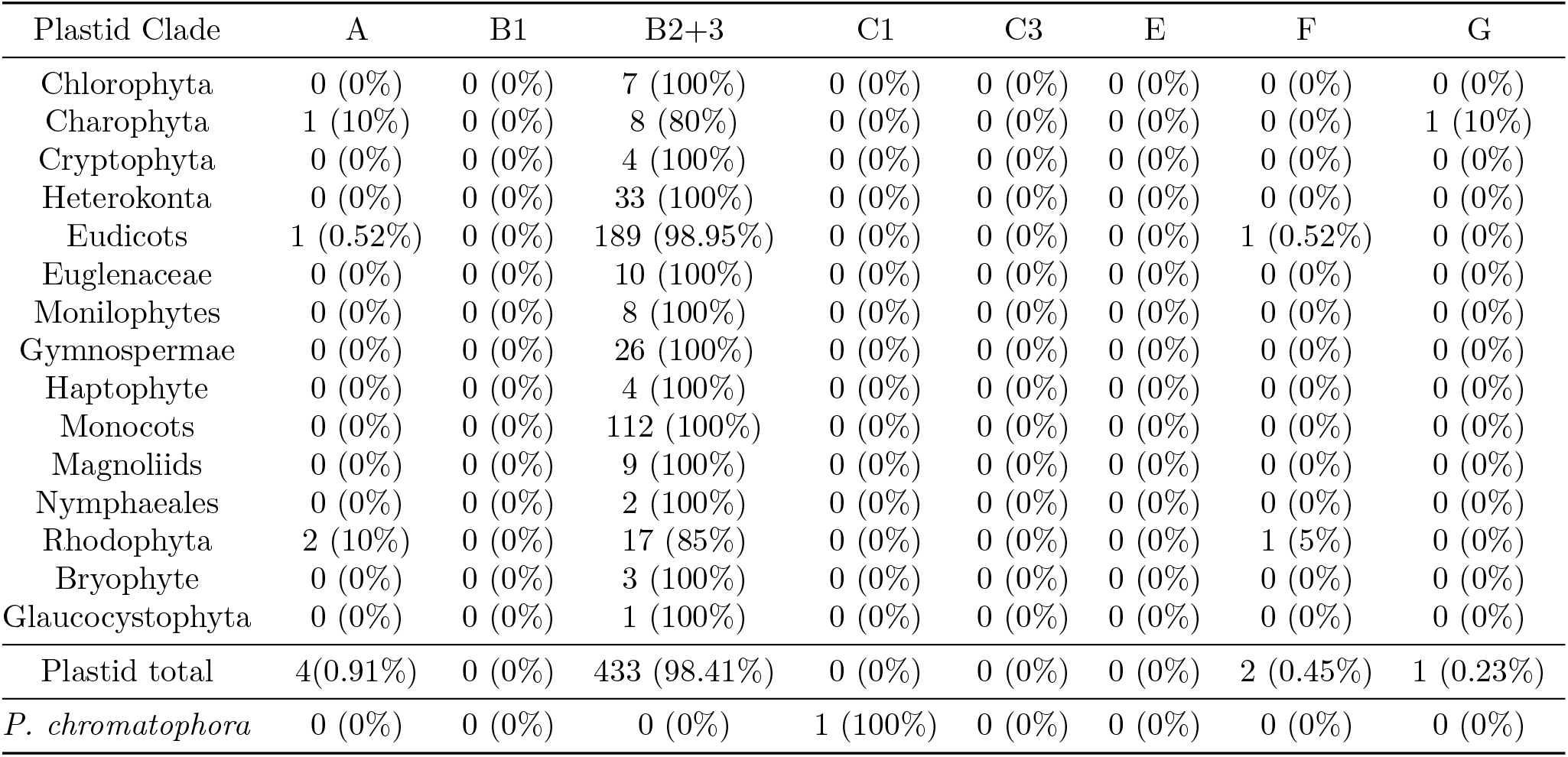
Classification results for plastid genomes and the chromatophore of *P. chromatophora* using CYANO-MLP. Results are summarized by plastid groups. Number of genomes classifying to each Cyanobacterial clade and percent are shown.

**Table 6.**
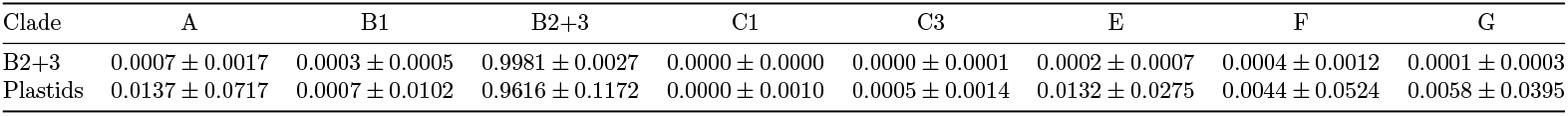
Mean probability plus and minus one standard deviation of classification of clade B2+3 and plastids to each Cyanobacterial clade using CYANO-MLP

**Table 7.**
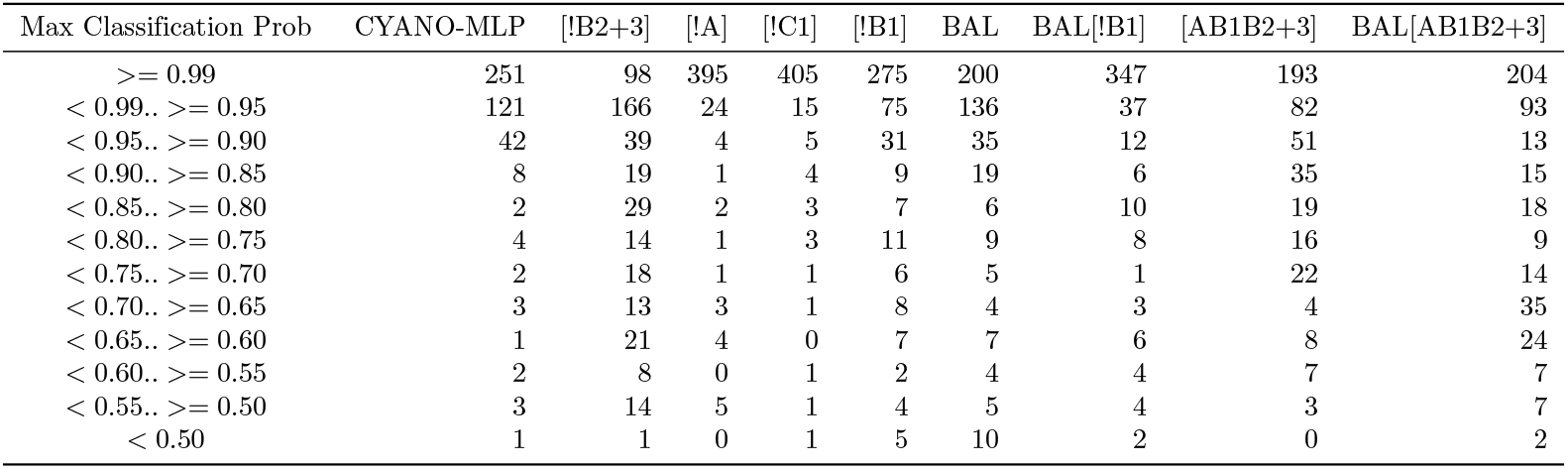
Number of plastid genomes with indicated max classification probability with the indicated version of CYANO-MLP

**Table 8.**
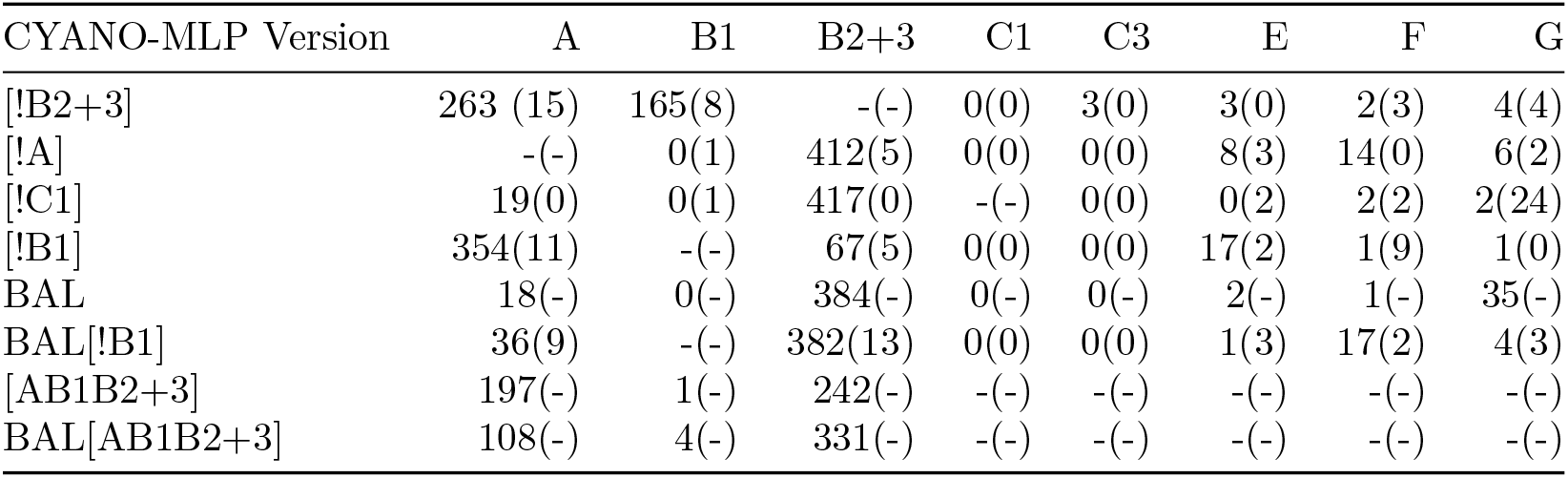
Number of plastid genomes and left-out cyanobacterial clade genomes classifying to each cyanobacterial clade for the indicated version of CYANO-MLP. The number outside of parentheses indicated number of plastid genomes and the number within parentheses is the left-out cyanobacterial clade genomes. Dashes indicate N/A values.

**Table 9.**
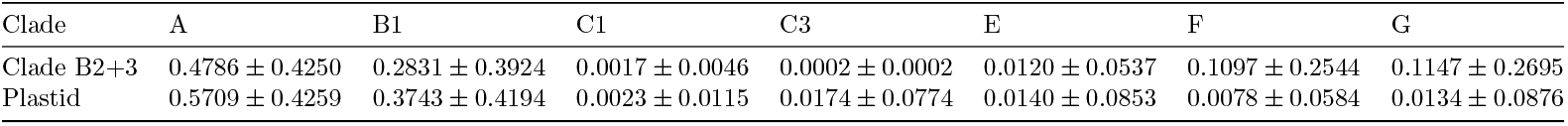
Mean probability and ± one standard deviation of classification for clade B2+3 and plastid genomes using CYANO-MLP[!B2+3].

**Table 10.**
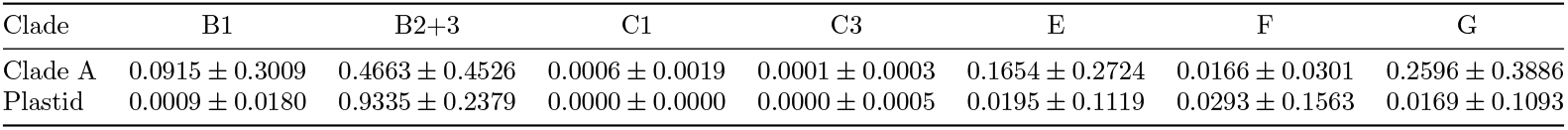
Mean probability and ± one standard deviation of classification for clade A and plastid genomes using CYANO-MLP[!A].

**Table 11.**
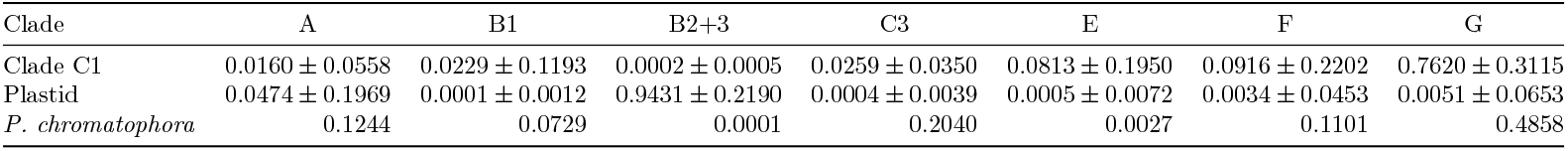
Mean probability and ± one standard deviation of classification for clade C1, plastid, and *P. chromatophora* chromatophore genome(s) using CYANO-MLP[!C1].

**Table 12.**
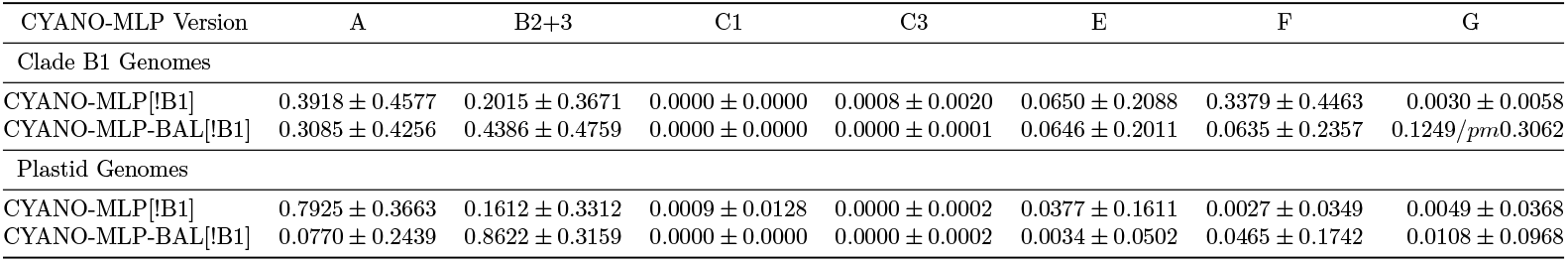
Mean probability ± one standard deviation of classification for B1 and plastid genomes using CYANO-MLP[!B1] and CYANO-MLP-BAL[!B1].

**Table 13.**
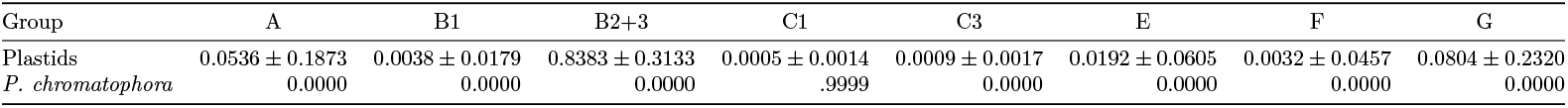
Mean probability of classification for plastid genomes and probability of classification for *P. chromatophora* using CYANO-MLP-BAL.

**Table 14.**
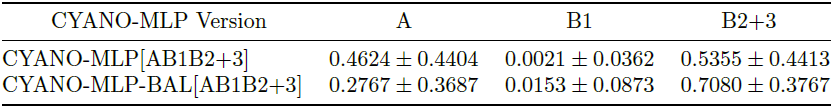
Mean probability ± one standard deviation of classification for plastid genomes using CYANO-MLP[AB1B2+3] and CYANO-MLP-BAL[AB1B2+3].

**Figure S21.**
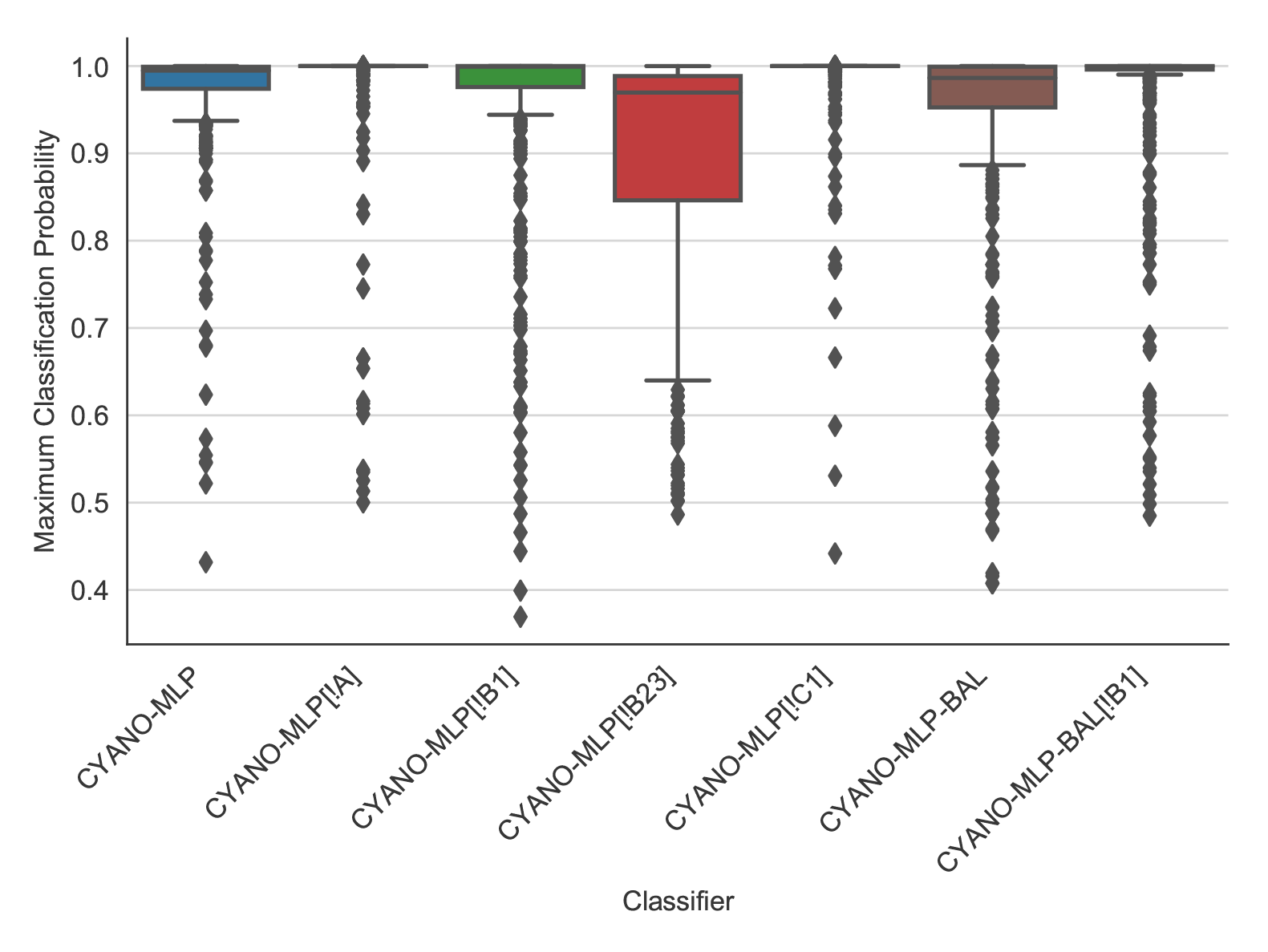
Box plot of maximum classification probability of each plastid genome. Error bars are the lesser of 1.5 IQR or full range of data.

**Table 15.**
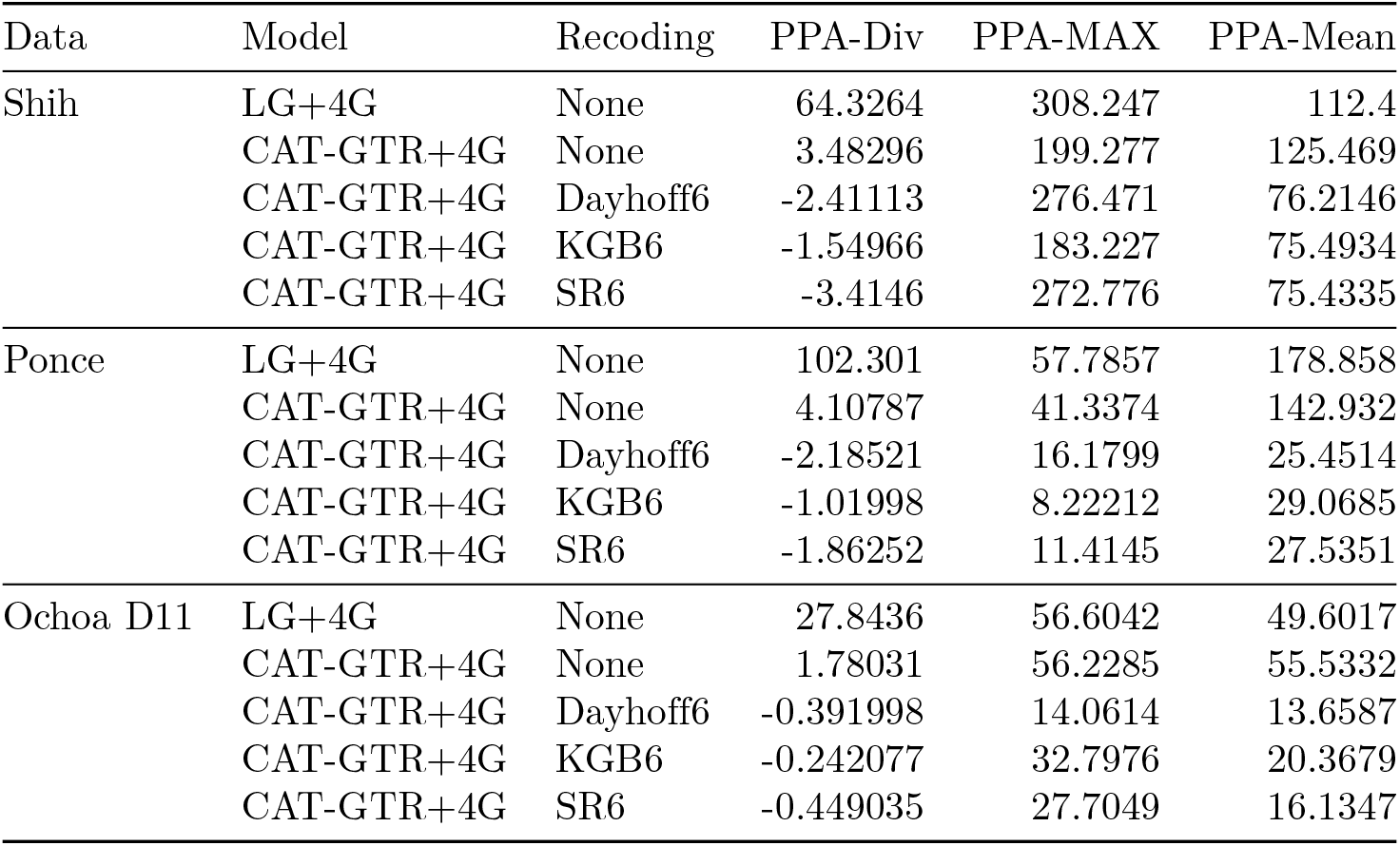
Results of the posterior predictive analyses presented as z-scores.

## Acknowledgments

DHA and TJL were supported by the National Science Foundation (INSPIRE-1344279). DHA was supported by NIH/NIAID 1R21AI127582-0. Computational research was performed on the MERCED HPC cluster supported by the National Science Foundation (ACI-1429783). The authors thank Harish Bhat, Emily Jane McTavish, Suzanne Sindi, Carolin Frank, Dana Carper, Jeanne Milostan and David Noelle for discussions.

## References

1. S. M. Adl, A. G. Simpson, C. E. Lane, J. Lukeš, D. Bass, S. S. Bowser, M. Brown, F. Burki, M. Dunthorn, V. Hampl, A. Heiss, M. Hoppenrath, E. Lara, L. leGall, D. H. Lynn, H. McManus, E. A. D. Mitchell, S. E. Mozley-Stanridge, L. W. Parfrey, J. Pawlowski, S. Rueckert, L. Shadwick, C. Schoch, A. Smirnov, and F. W. Spiegel. The revised classification of eukaryotes. Journal of Eukaryotic Microbiology, 59(5):429–493, 2012.

2. S. Alkatib, T. T. Fleischmann, L. B. Scharff, and R. Bock. Evolutionary constraints on the plastid tRNA set decoding methionine and isoleucine. Nucleic Acids Research, 40(14):6713–24, aug 2012.

3. K. C. H. Amrine, W. D. Swingley, and D. H. Ardell. tRNA signatures reveal a polyphyletic origin of SAR11 strains among alphaproteobacteria. PLoS Computational Biology, 10(2):e1003454, feb 2014.

4. D. H. Ardell and S. G. E. Andersson. TFAM detects co-evolution of tRNA identity rules with lateral transfer of histidyl-tRNA synthetase. Nucleic Acids Research, 34(3):893–904, 2006.

5. D. H. Ardell and Y.-M. Hou. Initiator tRNA genes template the 3′ CCA end at high frequencies in bacteria. BMC genomics, 17(1):1003, 2016.

6. S. Ball, C. Colleoni, U. Cenci, J. N. Raj, and C. Tirtiaux. The evolution of glycogen and starch metabolism in eukaryotes gives molecular clues to understand the establishment of plastid endosymbiosis. Journal of Experimental Botany, 62(6):1775–1801, mar 2011.

7. B. Batut, C. Knibbe, G. Marais, and V. Daubin. Reductive genome evolution at both ends of the bacterial population size spectrum. Nature Reviews Microbiology, 12(12):841, 2014.

8. C. E. Blank. Origin and early evolution of photosynthetic eukaryotes in freshwater environments: reinterpreting proterozoic paleobiology and biogeochemical processes in light of trait evolution. Journal of Phycology, 49(6):1040–1055, dec 2013.

9. C. E. Blank and P. Sánchez-Baracaldo. Timing of morphological and ecological innovations in the cyanobacteria–A key to understanding the rise in atmospheric oxygen. Geobiology, 8(1):1–23, jan 2010.

10. S. Blanquart and N. Lartillot. A Site- and Time-Heterogeneous Model of Amino Acid Replacement. Molecular Biology and Evolution, 25(5):842–858, may 2008.

11. T. Ching, D. S. Himmelstein, B. K. Beaulieu-Jones, A. A. Kalinin, B. T. Do, G. P. Way, E. Ferrero, P.-M. Agapow, M. Zietz, M. M. Hoffman, W. Xie, G. L. Rosen, B. J. Lengerich, J. Israeli, J. Lanchantin, S. Woloszynek, A. E. Carpenter, A. Shrikumar, J. Xu, E. M. Cofer, C. A. Lavender, S. C. Turaga, A. M. Alexandari, Z. Lu, D. J. Harris, D. DeCaprio, Y. Qi, A. Kundaje, Y. Peng, L. K. Wiley, M. H. S. Segler, S. M. Boca, S. J. Swamidass, A. Huang, A. Gitter, and C. S. Greene. Opportunities and obstacles for deep learning in biology and medicine. Journal of the Royal Society, Interface, 15(141):20170387, apr 2018.

12. A. Criscuolo and S. Gribaldo. Large-Scale Phylogenomic Analyses Indicate a Deep Origin of Primary Plastids within Cyanobacteria. Molecular Biology and Evolution, 28(11):3019, 2011.

13. T. Dagan, M. Roettger, K. Stucken, G. Landan, R. Koch, P. Major, S. B. Gould, V. V. Goremykin, R. Rippka, N. Tandeau de Marsac, M. Gugger, P. J. Lockhart, J. F. Allen, I. Brune, I. Maus, A. Pühler, and W. F. Martin. Genomes of Stigonematalean cyanobacteria (subsection V) and the evolution of oxygenic photosynthesis from prokaryotes to plastids. Genome biology and evolution, 5(1):31–44, 2013.

14. M. O. Dayhoff, R. M. Schwartz, and B. C. Orcutt. A model of evolutionary change in proteins. In Atlas of Protein Sequence and Structure, pages 345–352. 1978.

15. P. Deschamps, C. Colleoni, Y. Nakamura, E. Suzuki, J.-L. Putaux, A. Buleon, S. Haebel, G. Ritte, M. Steup, L. I. Falcon, D. Moreira, W. Loffelhardt, J. N. Raj, C. Plancke, C. D’Hulst, D. Dauvillee, and S. Ball. Metabolic Symbiosis and the Birth of the Plant Kingdom. Molecular Biology and Evolution, 25(3):536–548, jan 2008.

16. O. Deusch, G. Landan, M. Roettger, N. Gruenheit, K. V. Kowallik, J. F. Allen, W. Martin, and T. Dagan. Genes of Cyanobacterial Origin in Plant Nuclear Genomes Point to a Heterocyst-Forming Plastid Ancestor. Molecular Biology and Evolution, 25(4):748–761, feb 2008.

17. D. Domman, M. Horn, T. M. Embley, and T. A. Williams. Plastid establishment did not require a chlamydial partner. Nature communications, 6:6421, mar 2015.

18. A. Dufresne, M. Salanoubat, F. Partensky, F. Artiguenave, I. M. Axmann, V. Barbe, S. Duprat, M. Y. Galperin, E. V. Koonin, F. Le Gall, K. S. Makarova, M. Ostrowski, S. Oztas, C. Robert, I. B. Rogozin, D. J. Scanlan, N. Tandeau de Marsac, J. Weissenbach, P. Wincker, Y. I. Wolf, and W. R. Hess. Genome sequence of the cyanobacterium Prochlorococcus marinus SS120, a nearly minimal oxyphototrophic genome. Proceedings of the National Academy of Sciences of the United States of America, 100(17):10020–5, aug 2003.

19. S. R. Eddy and R. Durbin. RNA sequence analysis using covariance models. Nucleic Acids Research, 22(11):2079–2088, jun 1994.

20. L. I. Falcón, S. Magallón, and A. Castillo. Dating the cyanobacterial ancestor of the chloroplast. The ISME Journal, 4(6):777–783, jun 2010.

21. P. G. Foster and T. Schultz. Modeling Compositional Heterogeneity. Systematic Biology, 53(3):485–495, jun 2004.

22. E. Freyhult, V. Moulton, and D. H. Ardell. Visualizing bacterial tRNA identity determinants and antideterminants using function logos and inverse function logos. Nucleic Acids Research, 34(3):905–916, 2006.

23. J. Gorodkin, L. Heyer, S. Brunak, and G. Storomo. Displaying the information contents of structural RNA alignments: the structure logos. Bioinformatics, 13(6):583–586, dec 1997.

24. M. Gouy, S. Guindon, and O. Gascuel. SeaView Version 4: A Multiplatform Graphical User Interface for Sequence Alignment and Phylogenetic Tree Building. Molecular Biology and Evolution, 27(2):221–224, 2010.

25. P. J. Keeling. Diversity and evolutionary history of plastids and their hosts. American Journal of Botany, 91(10):1481–1493, oct 2004.

26. P. Kenrick and P. R. Crane. The origin and early evolution of plants on land. Nature, 389(6646):33–39, sep 1997.

27. C. Kosiol, N. Goldman, and N. H. Buttimore. A new criterion and method for amino acid classification. Journal of Theoretical Biology, 228(1):97–106, may 2004.

28. N. Lartillot, H. Brinkmann, and H. Philippe. Suppression of long-branch attraction artefacts in the animal phylogeny using a site-heterogeneous model. BMC Evolutionary Biology, 7(Suppl 1):S4, feb 2007.

29. N. Lartillot and H. Philippe. A Bayesian Mixture Model for Across-Site Heterogeneities in the Amino-Acid Replacement Process. Molecular Biology and Evolution, 21(6):1095–1109, jun 2004.

30. N. Lartillot, N. Rodrigue, D. Stubbs, and J. Richer. PhyloBayes MPI: Phylogenetic Reconstruction with Infinite Mixtures of Profiles in a Parallel Environment. Systematic Biology, 62(4):611–615, jul 2013.

31. D. Laslett and B. Canback. ARAGORN, a program to detect tRNA genes and tmRNA genes in nucleotide sequences. Nucleic Acids Research, 32(1):11–16, jan 2004.

32. T. J. Lawrence, K. T. Kauffman, K. C. Amrine, D. L. Carper, R. S. Lee, P. J. Becich, C. J. Canales, and D. H. Ardell. FAST: FAST Analysis of Sequences Toolbox. Frontiers in Genetics, 6, 2015.

33. S. Q. Le and O. Gascuel. An Improved General Amino Acid Replacement Matrix. Molecular Biology and Evolution, 25(7):1307–1320, apr 2008.

34. T. M. Lowe and S. R. Eddy. tRNAscan-SE: A Program for Improved Detection of Transfer RNA Genes in Genomic Sequence. Nucleic Acids Research, 25(5):0955–964, mar 1997.

35. J. R. Manhart and J. D. Palmer. The gain of two chloroplast tRNA introns marks the green algal ancestors of land plants. Nature, 345(6272):268–270, may 1990.

36. W. Martin and K. Kowallik. Annotated English translation of Mereschkowsky’s 1905 paper ‘Ü ber Natur und Ursprung der Chromatophoren im Pflanzenreiche’. European Journal of Phycology, 34(3):287–295, aug 1999.

37. G. I. McFadden and G. G. van Dooren. Evolution: red algal genome affirms a common origin of all plastids. Current biology, 14(13):R514–6, jul 2004.

38. C. Mereschkowsky. Über natur und ursprung der chromatophoren im pflanzenreiche. Biologisches Centralblatt, 25:593–604, 1905.

39. J. A. G. Ochoa de Alda, R. Esteban, M. Luz Diago, and J. Houmard. The plastid ancestor originated among one of the major cyanobacterial lineages. Nature Communications, 5:4937, sep 2014.

40. L. W. Parfrey, E. Barbero, E. Lasser, M. Dunthorn, D. Bhattacharya, D. J. Patterson, and L. A. Katz. Evaluating Support for the Current Classification of Eukaryotic Diversity. PLoS Genetics, 2(12):e220, dec 2006.

41. F. Pedregosa, G. Varoquaux, A. Gramfort, V. Michel, B. Thirion, O. Grisel, M. Blondel, P. Prettenhofer, R. Weiss, V. Dubourg, J. Vanderplas, A. Passos, D. Cournapeau, M. Brucher, M. Perrot, and E. Duchesnay. Scikit-learn: Machine learning in python. Journal of Machine Learning Research, 12:2825–2830, 2011.

42. H. Philippe and B. Roure. Difficult phylogenetic questions: more data, maybe; better methods, certainly. BMC Biology, 9(1):91, dec 2011.

43. R. I. Ponce-Toledo, P. Deschamps, P. López-García, Y. Zivanovic, K. Benzerara, and D. Moreira. An Early-Branching Freshwater Cyanobacterium at the Origin of Plastids. Current Biology, 27(3):386–391, feb 2017.

44. G. Rocap, F. W. Larimer, J. Lamerdin, S. Malfatti, P. Chain, N. A. Ahlgren, A. Arellano, M. Coleman, L. Hauser, W. R. Hess, Z. I. Johnson, M. Land, D. Lindell, A. F. Post, W. Regala, M. Shah, S. L. Shaw, C. Steglich, M. B. Sullivan, C. S. Ting, A. Tolonen, E. A. Webb, E. R. Zinser, and S. W. Chisholm. Genome divergence in two Prochlorococcus ecotypes reflects oceanic niche differentiation. Nature, 424(6952):1042–1047, aug 2003.

45. B. E. Schirrmeister, M. Gugger, and P. C. J. Donoghue. Cyanobacteria and the Great Oxidation Event: evidence from genes and fossils. Palaeontology, 58(5):769–785, sep 2015.

46. T. D. Schneider. A brief review of molecular information theory. Nano Communication Networks, 1(3):173 – 180, 2010. Fundamentals of Nanoscale Communications.

47. P. M. Shih, D. Wu, A. Latifi, S. D. Axen, D. P. Fewer, E. Talla, A. Calteau, F. Cai, N. Tandeau de Marsac, R. Rippka, M. Herdman, K. Sivonen, T. Coursin, T. Laurent, L. Goodwin, M. Nolan, K. W. Davenport, C. S. Han, E. M. Rubin, J. A. Eisen, T. Woyke, M. Gugger, and C. A. Kerfeld. Improving the coverage of the cyanobacterial phylum using diversity-driven genome sequencing. Proceedings of the National Academy of Sciences of the United States of America, 110(3):1053–8, jan 2013.

48. D. Simon, D. Fewer, T. Friedl, and D. Bhattacharya. Phylogeny and Self-Splicing Ability of the Plastid tRNA-Leu Group I Intron. Journal of Molecular Evolution, 57(6):710–720, dec 2003.

49. M. Sprinzl, C. Horn, M. Brown, A. Ioudovitch, and S. Steinberg. Compilation of tRNA sequences and sequences of tRNA genes. Nucleic acids research, 26(1):148–53, jan 1998.

50. M. Sugiura and T. Wakasugi. Compilation and comparison of transfer RNA genes from tobacco chloroplasts. Critical Reviews in Plant Sciences, 8(2):89–101, jan 1989.

51. E. Susko and A. J. Roger. On Reduced Amino Acid Alphabets for Phylogenetic Inference. Molecular Biology and Evolution, 24(9):2139–2150, may 2007.

52. P. Sánchez-Baracaldo, J. A. Raven, D. Pisani, and A. H. Knoll. Early photosynthetic eukaryotes inhabited low-salinity habitats. Proceedings of the National Academy of Sciences of the United States of America, 114(37):E7737–E7745, sep 2017.

53. J. N. Timmis, M. A. Ayliffe, C. Y. Huang, and W. Martin. Endosymbiotic gene transfer: organelle genomes forge eukaryotic chromosomes. Nature Reviews Genetics, 5(2):123–135, feb 2004.

54. J. Vogel, T. Börner, and W. R. Hess. Comparative analysis of splicing of the complete set of chloroplast group II introns in three higher plant mutants. Nucleic Acids Research, 27(19):3866–74, oct 1999.

